# A hybrid open-top light-sheet microscope for multi-scale imaging of cleared tissues

**DOI:** 10.1101/2020.05.06.081745

**Authors:** Adam K. Glaser, Kevin W. Bishop, Lindsey A. Barner, Etsuo A. Susaki, Shimpei I. Kubota, Gan Gao, Robert B. Serafin, Pooja Balaram, Emily Turschak, Philip R. Nicovich, Hoyin Lai, Luciano A.G. Lucas, Yating Yi, Eva K. Nichols, Hongyi Huang, Nicholas P. Reder, Jasmine J. Wilson, Ramya Sivakumar, Elya Shamskhou, Caleb R. Stoltzfus, Xing Wei, Andrew K. Hempton, Marko Pende, Prayag Murawala, Hans U. Dodt, Takato Imaizumi, Jay Shendure, Brian J. Beliveau, Michael Y. Gerner, Li Xin, Hu Zhao, Lawrence D. True, R. Clay Reid, Jayaram Chandrashekar, Hiroki R. Ueda, Karel Svoboda, Jonathan T.C. Liu

## Abstract

Light-sheet microscopy has emerged as the preferred means for high-throughput volumetric imaging of cleared tissues. However, there is a need for a user-friendly system that can address imaging applications with varied requirements in terms of resolution (mesoscopic to sub-micrometer), sample geometry (size, shape, and number), and compatibility with tissue-clearing protocols and sample holders of various refractive indices. We present a ‘hybrid’ system that combines a novel non-orthogonal dual-objective and conventional (orthogonal) open-top light-sheet architecture for versatile multi-scale volumetric imaging.

## Main text

Recent advances in tissue-clearing protocols greatly reduce optical scattering, aberrations, and background fluorescence, enabling deep-tissue imaging with high resolution and contrast. These approaches have yielded new insights in many fields, including neuroscience, developmental biology, and anatomic pathology [1-11]. Light-sheet microscopy has emerged as a preferred means for high-resolution volumetric imaging of cleared tissues due to its unrivaled speed and low photobleaching [12, 13]. Many variants of light-sheet microscopes have been developed in recent years by academic researchers and commercial entities to tackle a diverse range of imaging applications (Error! Reference source not found. and Error! Reference source not found.) [14-18]. Whereas individual light-sheet systems are well-suited for a subset of cleared-tissue applications, trade-offs are inevitable. In particular, no current light-sheet microscope can satisfy all of the following requirements: (1) user-friendly mounting of multiple specimens with standard holders, (2) compatibility with all current clearing protocols, (3) no fundamental limits on lateral specimen size, (4) a large imaging depth to accommodate intact mouse organs and thick tissue slabs, and (5) a wide range of optical resolutions for time/data-efficient imaging from sub-micrometer to mesoscopic scales (i.e., ‘multi-scale’ imaging). A system that meets these diverse requirements would satisfy a growing base of researchers interested in rapidly screening large intact cleared tissues followed by detailed interrogation of sub-micrometer structures within localized subregions.

The majority of cleared-tissue microscopes use an illumination objective that is oriented in the horizontal plane of the specimen, along with a collection objective that is orthogonal to the illumination objective, i.e., an orthogonal dual-objective (ODO) configuration [16, 19] (**Figure 1a**). This imaging geometry places physical constraints on the lateral size of the specimen such that it is not possible to image large centimeter-scale tissue slabs or multiple specimens mounted in standard holders (e.g., well plates). Therefore, we and others have previously developed ODO systems with an inverted geometry, which overcomes these constraints by tilting both objectives and placing them above or below the specimen [20-27] (**Figure 1b**). In particular, open-top light-sheet (OTLS) microscopy systems provide an ease-of-use similar to a flat-bed document scanner by placing all optical components below the sample holder [22-27]. This enables a wide range of modular sample holders to be used (e.g. well plates), as well as potential accessory technologies such as microfluidics, electrophysiology, and micro-dissection/aspiration. With OTLS systems, the angled orientation of the objectives reduces the usable imaging depth of the systems to well below each objective’s native working distance. In addition, in OTLS microscopy, angling the optical paths with respect to a horizontal sample holder can introduce significant off-axis aberrations (especially for higher-numerical-aperture beams) unless the refractive index of the specimen and the sample holder are exquisitely well-matched (**Supplementary Figure 1**). This can reduce the ease of use and versatility of high-numerical-aperture (high-NA) OTLS systems.

**Figure 1.**
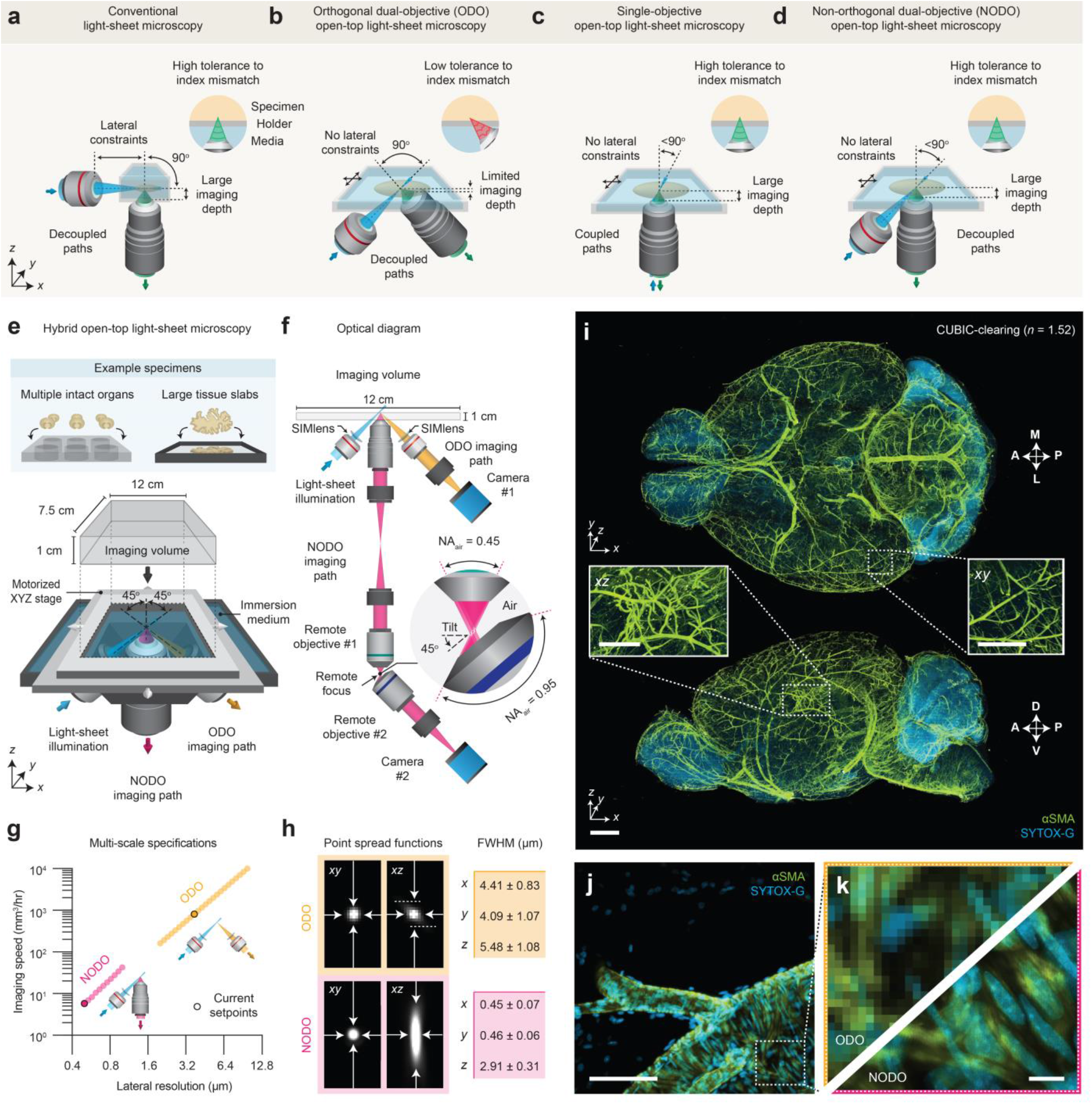
Hybrid open-top light-sheet (OTLS) microscopy. **(a)** Orthogonal dual-objective (ODO) light-sheet microscopes typically place one or more objectives in the horizontal plane of the specimen, which constrains the specimen size and prevents the use of standard modular sample holders. **(b)** ODO open-top light-sheet (OTLS) microscopy systems overcome this constraint by tilting the objectives and positioning them beneath a horizontal sample holder. However, a consequence of this is a limited imaging depth and low tolerance to refractive-index mismatch between the specimen and the sample-holder material (which can reduce ease-of-use). **(c)** Single-objective OTLS microscopy systems overcome both limitations. However, the use of a single objective for illumination and collection precludes broad multi-scale imaging and limits/couples the axial and lateral resolutions. **(d)** Non-orthogonal dual-objective (NODO) OTLS microscopy allows for laterally unconstrained imaging over the full working distance of a vertically oriented collection objective with high tolerance to index mismatch. The use of a separate objective for light-sheet illumination also enables more design flexibility to optimize axial and lateral resolutions. **(e)** The hybrid microscope architecture consists of three objectives, positioned below the specimen, mounted into a monolithic imaging chamber that is filled with an interchangeable immersion medium. One objective is used for light-sheet illumination, and the other two objectives are used for orthogonal dual-objective (ODO) and non-orthogonal dual-objective (NODO) collection. The ODO path provides fast meso-scale screening capabilities, and the NODO path enables targeted sub-micrometer imaging. By using a motorized stage, tiled imaging is possible with both paths over a large 12 × 7.5 × 1 cm (XYZ) imaging volume. This large imaging volume accommodates multiple intact cleared organs and large tissue slabs mounted in an array of specimen holders. The optical layout of the hybrid OTLS system is shown in **(f)**. The ODO path uses a single tube lens to directly image the light sheet onto Camera #1. The NODO path uses a remote focus system consisting of three objective-tube lens pairs to image the non-orthogonal light sheet onto Camera #2. **(g)** In combination, the ODO and NODO paths enable imaging over a tunable lateral resolution range of 0.5 – 10.7 μm at imaging speeds of ∼5 mm3 to 10 cm3 per hour. The current set points of the system are highlighted. **(h)** At these set points, the full-width half-maximum (FWHM) resolutions (*xyz*) for the ODO path are 4.09 ± 1.07, 4.41 ± 0.83, and 5.48 ± 1.08 μm, and for the NODO path are 0.45 ± 0.07, 0.46 ± 0.06, and 2.91 ± 0.31 μm (mean ± standard deviation). These values are measured from reflectance imaging of 150-nm gold beads in an ECi-cleared agarose phantom (638-nm illumination). **(i)** Representative ODO imaging results of an entire intact CUBIC-cleared mouse brain with arterial (αSMA) and nuclear (SYTOX-G) staining. The ODO imaging path, with near-isotropic resolution, is able to clearly resolve vasculature in both the *xy* and *xz* planes (insets). **(j-k)** Targeted imaging of a sub-region centered on a branching arteriole using the NODO imaging path resolves individual smooth muscle cells and sub-nuclear features that are not resolved by ODO imaging. Scale-bar lengths are as follows: **(i)** 1 mm (insets, 500 μm), **(j)** 100 μm, and **(k)** 10 μm. All images are displayed without deconvolution.

To overcome these issues, we considered the use of a single-objective architecture that has gained popularity in recent years, in which the illumination and collection beams share a single objective [28-40] (**Figure 1c**). For an open-top version of single-objective light-sheet microscopy, orienting the objective in the vertical direction with respect to a horizontal specimen holder makes use of the objective’s full working distance and dramatically increases the system’s tolerance to refractive-index mismatch (**Supplementary Figure 2**). However, single-objective OTLS microscopy is incompatible with multi-scale imaging because, to the best of our knowledge, no currently available objective provides both sub-micrometer resolution and a large mesoscopic field of view (FOV). In addition, the use of a single objective both constrains and couples the illumination and collection beams such that there is less flexibility to tailor the axial and lateral resolutions (**Supplementary Figure 3**) [41]. Recently, we have explored the concept of a non-orthogonal dual-objective (NODO) OTLS configuration that makes use of a high-NA collection objective oriented in the vertical direction (similar to single-objective OTLS microscopy) in conjunction with a separate angled objective to provide lower-NA light-sheet illumination (**Figure 1d**). Our simulations indicate that a NODO configuration maintains the high index-mismatch tolerance of single-objective OTLS microscopy (**Supplementary Figure 4**) and offers increased flexibility to optimize lateral and axial resolutions [41].

To address the varied requirements of cleared-tissue light-sheet microscopists, we designed a ‘hybrid’ OTLS microscope that is the first system to use a non-orthogonal dual-objective (NODO) configuration (for high-resolution imaging). For low-resolution imaging, a conventional ODO open-top system is integrated with the NODO system. This new hybrid system leverages the strengths and overcomes the limitations of previous systems (**Supplementary Note 1**), addressing the five requirements listed earlier: (1) simple open-top mounting of multiple specimens in standard holders such as well plates, (2) the ability to pair any tissue-clearing reagent with nearly any sample-holder material with negligible degradation in imaging performance (i.e., high tolerance to refractive-index mismatch, **Supplementary Figure 5**), (3) no lateral constraints on the specimen size (limited only by the travel range of the microscope stage), (4) a 1-cm imaging depth for comprehensive interrogation of intact mouse organs and thick tissue slabs, and (5) multi-scale imaging over an unprecedented range roughly corresponding to what is achieved with 2X to 40X objectives. These unique capabilities open the door for new light-sheet microscopy applications, including efficient multi-scale imaging workflows in which one or more large specimens must be rapidly screened at low resolution to identify localized regions of interest for quantitative interrogation at the sub-micrometer scale.

The layout of our new system is shown in **Figure 1e**. The system architecture features three main objectives that are selected to avoid geometric interference. All three objectives are positioned below the specimen, which provides an unobstructed open top that enables volumetric imaging over a large 12 × 7.5 × 1 cm (*xyz*) imaging volume. The imaging volume is limited in *z* by the objectives and in *xy* by the mechanical limits of the motorized stage. All three objectives are sealed into a monolithic imaging chamber through direct immersion or the use of a solid-immersion meniscus lens (SIMlens), which provides multi-immersion capabilities spanning the refractive index range of all current clearing protocols (**Supplementary Figure 6**) [26, 27].

The optical layout of the system is shown in **Figure 1f**. To achieve an optimal combination of sub-micrometer resolution, large imaging depth, and compatibility with standard sample holders, we developed a new NODO light-sheet configuration [41]. By using the full numerical aperture (NA) of a vertically oriented objective for fluorescence collection, and a separate objective for non-orthogonal illumination, our new NODO architecture provides superior resolution to that of a single-objective light-sheet system (if based on the same primary objective) and relaxes the NA requirements for the remote-focus module that is necessary for these non-orthogonal light-sheet systems (**Supplementary Figure 4**). This allows for the use of a wider range of moderate-NA primary (collection) objectives for cleared-tissue imaging, as well as simple air objectives at the remote-focus module rather than bespoke objective assemblies (**Supplementary Figure 7**). We carefully selected our NODO collection objective and its optical and mechanical specifications to enable the placement of a separate low-NA collection objective oriented orthogonally to the illumination objective **(Supplementary Figure 8)**. This forms an additional ODO imaging path for rapid mesoscopic-resolution imaging. Similar to single-objective light-sheet systems, the NODO path uses a remote focus to re-image the non-orthogonal light sheet onto a camera, whereas the ODO collection path directly images the orthogonal light sheet onto a camera (**Supplementary Figures 9 – 16** and **Supplementary Video 1** contain optical models, a photograph of the hybrid OTLS microscope, and additional point spread functions for both imaging paths). The same illumination path is used to generate a light sheet with a variable width for both the NODO and ODO imaging paths.

The overall magnification of each imaging path is tunable, limited at the extremes by either the NA or FOV of the collection objectives (see **Supplementary Note 2** for further discussion of these two imaging modes), corresponding to a combined lateral-resolution range of ∼0.4 – 10 μm, and volumetric imaging speeds of ∼5 mm^3^ to 10 cm^3^ per hour (**Figure 1g)**. In the current microscope configuration, the NODO path provides an XYZ resolution of 0.45 ± 0.07, 0.46 ± 0.05, and 2.91 ± 0.31 μm (*N* = 437 beads), and the ODO path provides an XYZ resolution of 4.41 ± 0.83, 4.09 ± 1.07, and 5.48 ± 1.08 μm (*N* = 109 beads) (mean ± standard deviation, **Figure 1h**). Representative imaging results of a CUBIC-cleared mouse brain labeled with brain-wide arterial (αSMA) and nuclear (SYTOX-G) stains are shown in **Figures 1i-k** (**Supplementary Video 2**). The entire intact specimen is rapidly imaged with near-isotropic resolution using the mesoscopic ODO path of the hybrid system, which clearly resolves vasculature in all three dimensions. Detailed interrogation of a sub-region with the high-resolution NODO path resolves individual smooth muscle cells and sub-nuclear features, which are not resolved in ODO images of the same sub-region. We further spotlight the unique utility of our hybrid system in two example applications where multi-scale imaging enables time- and data-efficient experimental workflows.

First, we imaged axons in an intact mouse brain cleared in ethyl cinnamate (ECi) (**Supplementary Video 3**). Tracking the axons of individual neurons is a challenging problem – axons can be very thin (100 nm) and span very large distances (cm). To do this effectively, one typically relies on sparse and bright labeling of a few neurons along with high-resolution, high-contrast imaging of the entire brain (∼0.5 cm^3^) [42, 43]. Sub-micrometer imaging of such large volumes generates data sets that are tens of terabytes in size, necessitating computationally intensive downstream pipelines for data handling, processing, and storage. The multi-scale imaging capability of our hybrid system greatly accelerates and simplifies this process by screening an entire brain at low resolution to identify target regions with imageable neurons, followed by high-resolution imaging of the identified regions of interest.

Using the ODO imaging path, a mouse brain was rapidly screened in ∼1 hour with isotropic ∼2-μm voxels (∼4-μm resolution), revealing brain-wide axonal projections (**Figures 2a-b**). Inspection of dense axonal projections in *xy* and *xz* planes through the midbrain confirms the ability of the system to provide near-isotropic 4-to 5-μm resolution throughout the entire intact brain (**Figure 2c**). A targeted region of interest around a cortical pyramidal neuron was then imaged at <0.5-um *xy* resolution using the NODO imaging path (**Figure 2d-e**). This imaging resolution is sufficient to discern spines and varicosities on individual dendrites and axons (**Figures 2f-h**). Importantly, imaging a 1-mm^3^ subregion containing the targeted neuron at ∼0.5 × 0.5 × 2.7 μm (*xyz*) resolution required only 24 minutes, generating a 180-GB data set. By comparison, imaging the entire 0.5-cm^3^ brain volume at this resolution would require ∼2 weeks of imaging and would produce 200 TB of imaging data.

**Figure 2.**
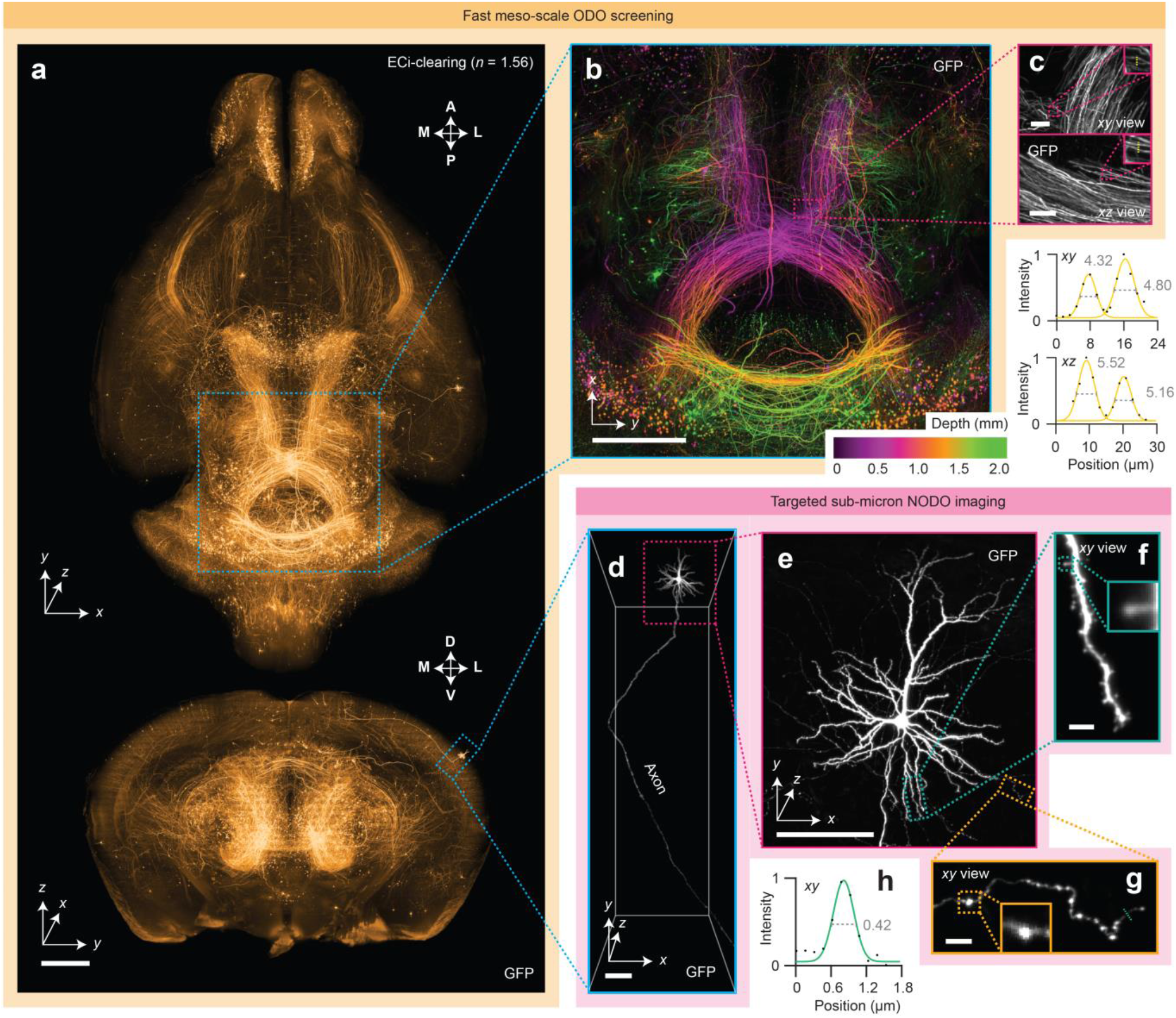
Fast meso-scale screening and targeted sub-micrometer imaging in cleared tissues. **(a)** Fast meso-scale screening is performed of an entire intact ECi-cleared Slc17a7-Cre mouse brain with brain-wide axonal projections. **(b)** A depth-coded region of interest shows dense projections in the midbrain. **(c)** *xy* and *xz* zoom-in views illustrate the near-isotropic resolution of the hybrid OTLS microscope. Line profiles through individual axons demonstrate an ODO lateral and axial resolution of 4-5 μm at a large depth within the cleared specimen. **(d-e)** Targeted sub-micrometer imaging is performed of a region of interest around a cortical pyramidal neuron. **(f-g)** Zoom-ins of a dendrite and axon demonstrate sufficient lateral resolution to visualize individual spines and varicosities. **(h)** A line profile through an individual axon demonstrates a NODO lateral resolution of 0.42 μm within the cleared specimen. Scale-bar lengths are as follows: **(a-b)** 1 mm, **(c)** 10 μm, **(d-e)** 100 μm, and **(f-g)** 5 μm. All images are displayed without deconvolution.

In a second example of the unique capabilities of our hybrid OTLS system, an imaging experiment was performed that required broad multi-scale imaging across multiple specimens. We used our hybrid OTLS system to study metastatic colonies from two cancer cell lines (MDA-231 and OS-RC-2) throughout intact mouse brains (**Supplementary Video 4**). Due to the sparse and unpredictable spatial distribution of brain metastases, identifying those sites in whole brains is challenging without a rapid low-resolution screening method. Once metastatic sites are identified, high-resolution quantitative analysis of these regions is also desired. While past studies have relied on laborious and time-consuming experiments with manual transfer of individual specimens between different microscope systems [44] (for low-resolution localization followed by high-resolution analysis), we analyzed six intact mouse brains in a single imaging session without manually removing or remounting the specimens (**Figure 3a**).

**Figure 3.**
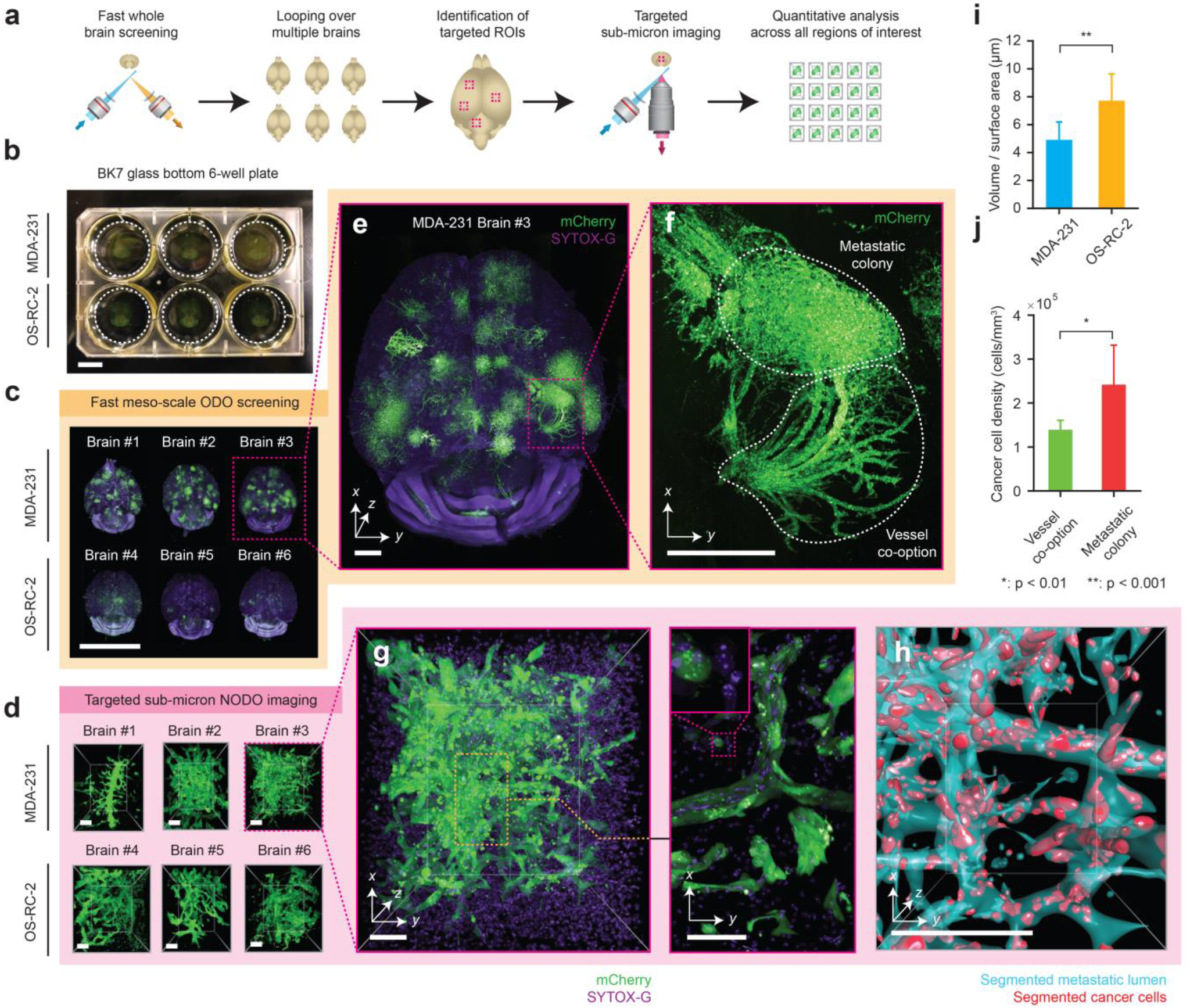
Multi-scale OTLS microscopy for quantitative analysis of brain metastases. **(a)** Hybrid OTLS microscopy workflow for multi-scale imaging of multiple specimens. **(b)** *N* = 6 whole mouse brains containing metastatic lesions from two different cancer cell lines (MDA-231 and OS-RC-2) were placed into a glass-bottom 6-well plate and mounted onto the hybrid OTLS microscope system for multi-scale imaging. **(c)** The ODO imaging path was used to rapidly screen multiple intact mouse brains containing metastatic colonies. **(d)** *N* = 34 total metastatic regions of interest (ROI) across all brains were identified and subsequently imaged at sub-micrometer resolution using the NODO imaging path (only one ROI per brain is shown for illustrative purposes). Visual inspection of a single brain with MDA-231 metastases in **(e)** revealed multiple colonies distributed throughout the brain, with signs of vessel co-option **(f-g)** that were not observed for OS-RC-2 metastases. **(h)** To quantify these phenotypic differences between the MDA-231 and OS-RC-2 metastases, the metastatic lumens (cyan) and cancer cells (red) were computationally segmented in all *N* = 34 ROIs. Quantification of the resulting segmentation masks revealed statistically (two-sample t-test) different 3D growth patterns between the two cancer cell lines, consistent with a previous report **(i-j)** [44]. Error bars in **(i-j)** denote standard deviation. Scale-bar lengths are as follows: **(b-c)** 1 cm, **(d)** 100 μm, **(e)** 1 mm, **(f)** 500 μm, and **(g-h)** 100 μm. All images are displayed without deconvolution.

All mouse brains were first placed in a standard 6-well plate (**Figure 3b**) and screened sequentially using fast mesoscopic ODO imaging (**Figure 3c**). After imaging, 34 total regions of interest (ROI) centered on metastatic colonies were manually identified across all mouse brains and sequentially imaged without having to move the specimens or specimen holder. An example ROI from each brain is shown in **Figure 3d**, with all 34 ROIs shown in **Supplementary Figure 17**. Visual inspection of whole-brain images reveals that metastatic growth from MDA-231 cancer cells exhibit vessel co-option (**Figures 3e-g)** whereas OS-RC-2 cancer-cell metastases do not. Morphological analysis of volumetrically segmented brain metastases, performed across the high-resolution ROIs, reveal statistically significant differences between the two metastatic cell lines. For example, the “volume per surface area” shape metric, which correlates with the roundness of a metastatic colony, was found to be higher for the OS-RC-2 cell line (**Figure 3i**). In addition, for the MDA-231 cancer cell line, the density of cancer cells in the vessel co-option ROIs was found to be lower than the adjacent metastatic colonies (**Figure 3j**). Both of these findings are consistent with the previous report from Kubota et al [44]. However, while this previous study required two separate microscope systems and several weeks of tedious imaging, including manual specimen transfer and co-registration of coordinates (personal correspondence with S.I.K.), our hybrid OTLS system allowed the entire imaging experiment to be completed in ∼1 day (∼2 hours of low-resolution imaging time per whole brain and ∼20 minutes of high-resolution imaging time per ROI).

Compared to existing academic and commercial light-sheet microscopy systems, our hybrid OTLS system provides a unique combination of versatility and performance necessary to satisfy the diverse requirements of a growing number of cleared-tissue imaging applications. In addition to the imaging examples in **Figures 1-3**, the system has potential utility for multi-scale non-destructive 3D pathology of prostate cancer with ECi clearing (**Supplementary Figure 18**), whole brain imaging of endogenous fluorescent proteins with PEGASOS clearing (**Supplementary Figure 19 and Supplementary Video 5**), 3D imaging of mouse embryos with SHIELD clearing (**Supplementary Figure 20**), mapping of immune cell populations in 3D with Ce3D clearing (**Supplementary Figure 21 and Supplementary Video 6**), assessment of 3D cell proliferation with iDISCO clearing (**Supplementary Figure 22**), imaging of plants/pigmented animal models processed with ClearSee/DEEP-Clear (**Supplementary Figure 23**), screening of immunofluorescent and endogenously fluorescent mouse brains with CUBIC clearing (**Supplementary Figure 24 Supplementary Videos 7 - 9**), and large-scale imaging of amyloid plaques in thick human brain slices measuring up to 12 × 7.5 × 1 cm in size (**Supplementary Figure 25 and Supplementary Video 10**).

In the future, both the axial and lateral resolution of the hybrid OTLS system can be improved. For example, the axial resolution of the system can be enhanced by incorporating a Bessel beam or by using a higher-NA Gaussian beam for light-sheet illumination [17, 18, 45, 46] (**Supplementary Figure 26**). This is facilitated by the fact that unlike single-objective light-sheet systems, our new NODO architecture allows the illumination NA to be substantially increased without necessitating a concomitant decrease in the collection NA (i.e. decoupled optical paths as shown in **Figure 1d**). With the 45-deg orientation of the two low-NA objectives in our system (one on either side of the NODO collection objective), it may also be possible to perform dual-sided illumination with fusion deconvolution to achieve improved image quality (**Supplementary Figure 27**) [20, 37, 47]. The lateral resolution of the system can also be improved by increasing the collection NA of the multi-immersion objective. However, the optical cone angle cannot be increased beyond 40 – 45 deg for a hybrid NODO / ODO system, as this would prevent the ODO objectives from being positioned at 45 deg with respect to the vertical axis.

While our current design uses three separate objectives, it is theoretically possible to achieve improved performance with an optimized NODO objective (**Supplementary Figure 28**), or to achieve similar multi-scale OTLS performance using a single high-NA objective with a large FOV and working distance (see **Supplementary Note 4** and **Supplementary Figure 29**). For example, the pupil of a single objective could be split into three separate regions, using the edges for NODO illumination and for mesoscopic ODO imaging, and only the center for high-resolution NODO collection. However, this would require an objective with extraordinary specifications that could be prohibitively expensive to design and manufacture. Therefore, in summary, our hybrid OTLS design represents a practical means, with commercially available optical components, of achieving an impressive balance of performance and versatility for a growing number of cleared-tissue imaging experiments in which rapid low-resolution screening of large volumes is desired in addition to high-resolution characterization of localized subregions.

## Online Methods

### Hybrid open-top light-sheet microscope

Light-sheet-based imaging is achieved using three optical arms. The first obliquely illuminates the specimen with a light sheet at 45 deg relative to the vertical axis. The second is oriented vertically, enabling high-resolution collection of the light-sheet-generated fluorescence in a NODO configuration. Finally, the third arm is oriented at 90 deg relative to the illumination light sheet, enabling high-speed mesoscopic imaging in an ODO configuration. All three objectives are positioned below the specimen and are sealed into a custom-designed monolithic imaging chamber (machined by Hilltop Technologies). In the case of each angled air objective, a custom-fabricated solid-immersion meniscus lens (SIMlens) (fabricated by BMV Optical Technologies) is sealed into the chamber (**Supplementary Figure 6**) [26]. Stage-scanning is achieved by using an XY stage (MS2000, Applied Scientific Instrumentation) attached to two Z-axis risers (LS50, Applied Scientific Instrumentation). 405, 488, 561, and 638 nm excitation laser light is provided to the illumination path by a multi-line laser package (Cobolt Skyra, HÜBNER Photonics). The NODO collection path is equipped with a motorized filter wheel (FW102C, Thorlabs) with four filters that can be used with 405-nm (FF02-447/60-25, Semrock), 488-nm (FF03-525/50-25, Semrock), 561-nm (FF01-618/50-25, Semrock), or 638-nm (BLP01-647R-25, Semrock) excitation. The ODO collection path is equipped with a single multi-band-pass filter (FF01-432/515/595/730-25, Semrock) that can be used with any of the four excitation wavelengths. The entire system is compact and fits on a portable 2’ × 3’ optical cart (POC001, Thorlabs).

### Illumination optical path

The illumination optical arm is shown in **Supplementary Figure 9**. The illumination optics are designed to allow for the light-sheet properties (i.e., width, thickness, and depth of focus) to be adjusted. In addition, the optical path is designed to be compatible with the refractive index, *n*, of all current clearing protocols (i.e., it has multi-immersion capabilities) and to minimize chromatic aberrations and defocusing (i.e., variations in the illumination focal length as a function of wavelength).

Illumination light is fiber coupled into the system with a Gaussian numerical aperture (NA) of ∼0.12 and collimated using an objective (RMS20X, Olympus). The beam diameter is then adjusted using a 4X variable beam expander (BE052-A, Thorlabs). This serves to adjust the overall NA of the light sheet. The variably expanded beam is then passed through an electronically tunable lens (ETL) that enables axial adjustment/alignment of the light sheet (EL-16-40-TC-VIS-5D-C, Optotune). The axially adjusted beam is relayed 1:1 using a pair of lenses (AC254-75A, Thorlabs) so that it can be scanned using a pair of large-beam-diameter galvanometric scanning mirrors (GVS012, Thorlabs). One mirror is scanned to create a digitally scanned light sheet [48]. The other scanning mirror is used to align the light sheet with the focal plane of the NODO and ODO imaging paths. A pair of achromatic doublet lenses (AC508-75A and AC508-200A, Thorlabs) are then used to relay the scanned beam to the back focal plane of a 2X illumination objective with NA = 0.10 (TL2X-SAP, Thorlabs). Finally, the illumination light travels through the SIMlens (**Supplementary Figure 6**) [26]. The SIMlens provides multi-immersion performance and prevents aberrations (spherical, off-axis, and chromatic) of the light sheet by minimizing refraction of the illumination rays as they transition from air into the immersion medium. In addition, since ray angles are preserved as they transition between air and the immersion medium, the SIMlens increases the NA of the illumination light sheet by a factor of *n*. When combined, this optical design yields a light sheet with tunable NA (0.025 – 0.10 × *n*) and tunable width (0 – 11 mm / *n*) limited by the field of view (FOV) of the illumination objective.

A digitally scanned light sheet was chosen over a cylindrical-lens approach (static light sheet), as this facilitates achieving a high level of tunability for multi-scale imaging. Moreover, large scanning mirrors were selected to fill the back focal plane of the final illumination objective and to avoid having to significantly magnify the beam after the scanning mirrors, which would reduce the lateral scanning range of the mirrors (i.e., constrain the maximum light-sheet width). In the current design, rotating the scanning mirror results in lateral scanning at a ratio of ∼0.60 mm per deg / *n*. The maximum desired light sheet width (i.e., lateral scanning range) is 11 mm, corresponding to a scanning angle of ∼18 deg, which is within the maximum scan range of the galvo scanner (20 deg).

### NODO optical path

The physical layout, ZEMAX model, and objective options for this NA-maximized NODO imaging configuration are shown in **Supplementary Figure 10**. The NODO optical path of our system uses a multi-immersion objective (#54-12-8, Special Optics) with a long 1-cm working distance. This objective is compatible with all clearing protocols (*n* = 1.33 – 1.56) and provides a NA of 0.483 (in air) that scales with the index of the immersion medium (e.g., NA ∼ 0.75 at *n* = 1.56). The lens is oriented in the normal (vertical) direction with respect to the specimen holder/interface and is therefore non-orthogonal to the light sheet. To image this non-orthogonal light sheet, we use a remote-focus imaging strategy analogous to what is used for single-objective light-sheet systems, with the multi-immersion objective serving as the primary objective (O1) [28, 29].

To minimize aberrations in the remote focus relay, the overall magnification from the specimen to the remote focus (air) should be equal to the refractive index of the specimen, *n*. Given this requirement, the relay lenses and first remote objective (O2) must be carefully selected. For O2, a 20X objective is optimal, as a 10X or 40X objective would clip either the NA or FOV of our O1. Of the several companies that produce microscope objectives, Zeiss and Leica were avoided because chromatic aberrations are partially corrected in the tube lenses produced by these companies, which would complicate selection of the two relay tube lenses. Therefore, only objectives from Olympus and Nikon were considered, where 20X objectives from Olympus have a focal length of 9 mm, and 20X objectives from Nikon have a focal length of 10 mm. Factoring in the effective focal length of O1 (12.19 mm / *n*), the required relay lens magnification is ∼1.219X for Nikon and ∼1.354X for Olympus. Note that the magnification of the multi-immersion objective in our system scales inversely as a function of *n*. This allows us to use a fixed set of relay lenses and always satisfy the remote focusing magnification requirement. However, this would be problematic with alternative multi-immersion objectives, where the effective focal length does not vary with *n*, and therefore the magnification of the relay lenses would need to be adapted for each immersion medium.

Well-corrected tube lenses are available with a limited selection of focal lengths (100, 165, 180, and 200 mm). Although custom tube lens assemblies are possible [33], we found that off-the-shelf 200-mm (Nikon Tube Lens #58-520, Edmund Optics) and 165-mm (TTL165-A, Thorlabs) tube lenses provide a magnification of ∼1.212X, which matches the requirement for Nikon. Therefore, we decided to select a 20X Nikon objective for O2. To avoid the need for a cover glass, we narrowed our selection to Nikon objectives designed for use without a cover glass. This yielded one option, the LU (now TU) Plan Fluor EPI 20X (NA = 0.45). We chose this objective for the O2 in the current system.

To remotely correct for spherical aberrations introduced by the index mismatch of the specimen holder, it would also be possible to use a Nikon objective with a correction collar. There are two options, the CFI S Plan Fluor ELWD 20XC (NA = 0.45) with a correction collar for a cover glass of *t* = 0 – 2 mm, and CFI S Plan Fluor LWD 20XC (NA = 0.70) with a correction collar for a cover glass of *t* = 0 – 1.8 mm. Both objectives could be explored in a future design (although spherical aberrations were not found to be an issue in the current design). In the case of the CFI S Plan Fluor LWD 20XC, the full 0.483 NA of our O1 objective could be transmitted to the remote focus, unlike the chosen LU Plan Fluor EPI 20X objective or CFI S Plan Fluor ELWD 20XC objective, which both clip things down to 0.45 NA.

The goal of O3 is to maximize light collection when tilted at the angle required to orthogonally image a remote version of the oblique light sheet within the specimen. As mentioned previously, in a single-objective light-sheet design, the light sheet angle would be limited by our chosen O1 to a maximum of 28.9 deg. This would require O3 to be tilted by at least 61.1 deg. At this extreme tilt angle, one way to prevent light loss is through a custom solid- or liquid-immersion objective [32, 33]. However, a benefit of our NODO design is that the crossing angle can be increased to 45 deg, which reduces the tilt of O3 to 45 deg. At this tilt angle, an objective with NA = 0.95 is able to capture light up to NA = 0.45 and provide NA-maximized imaging. Therefore, we opted to use a NA = 0.95 air objective for our O3. This use of a tertiary air objective makes alignment more straightforward and stable than in the case of a solid- or liquid-immersion objective [32, 33]. This is especially the case for liquid-immersion objectives, where there may be evaporation or leakage of a liquid medium over time.

In terms of selecting an optimal O3 for NA-maximized imaging, we only considered objectives from Olympus and Nikon for the same reasons as mentioned previously. Both companies offer two types of objectives with NA = 0.95. These include 40X life-science objectives with a correction collar for cover-glass thicknesses ranging from *t* = 0.11 – 0.23 mm, and 50X metrology objectives for imaging with no cover glass. Although it would be possible to permanently align or adhere a cover glass to the 40X objectives, we decided to select the CF IC EPI Plan Apo 50X objective from Nikon for our O3 for simplicity and ease-of-use. When combined with a 100-mm tube lens (TTL100-A, Thorlabs), our NODO imaging path provides a total magnification of 25 × *n*. This yields a sampling rate of ∼2.71 (slightly better than Nyquist) when using a sCMOS camera with pixels spaced by 6.5 μm (pco.edge 4.1, PCO Tech). The corresponding FOV is ∼ 0.53 mm / *n* (FOV = 0.40 - 0.34 mm when *n* = 1.33 – 1.56), which is not clipped by the 0.40-mm FOV of the 50X objective. In this configuration, the back aperture of the illumination objective is filled, yielding an illumination NA of ∼0.10 × *n*. This corresponds to a confocal parameter of ∼40 – 50 μm and a pixel height of ∼256 pixels for each raw camera image. This also corresponds to an axial re-focusing range that is well within the range of operation for an idealized objective, as specified by Botcherby et al. [28]. The pixel width of each raw camera frame is the full 2048 pixels of the sCMOS camera, corresponding to a FOV of ∼0.34 – 0.40 mm.

While the above O3 selection provides NA-maximized imaging, our O1 also provides a 1-mm FOV to enable FOV-maximized imaging (with slightly worse resolution). To achieve this, O3 can be changed to a 20X objective with a matched 1-mm FOV. To minimize light loss, 20X objectives with the highest NA were considered. For both Olympus and Nikon, the upper limit is NA = 0.75 (although the recent UPLXAPO20X objective from Olympus offers NA = 0.80). In a single-objective light-sheet configuration, the light sheet angle in the specimen would be limited by the NA of O1, requiring a larger remote tilt angle, which for this particular O3 would result in an effective NA ∼ 0.12 × *n*. In contrast, with our NODO configuration this results in an effective NA ∼ 0.26 × *n*. This reduced effective NA (in contrast to the NA-maximized case) enables close-to-Nyquist sampling of a 1-mm FOV with the same sCMOS chip as described earlier (with 2048 × 2048 pixels). To match parfocal lengths and thus enable a quick interchange of objectives, we selected the Nikon CFI Plan Apo Lambda 20X for our FOV-maximized O3 objective. When paired with the same 100-mm tube lens (TTL100-A, Thorlabs), this provides a total magnification of 10 × *n*, which yields a near-Nyquist sampling rate of 2.02. The effective FOV is ∼ 1.33 mm / *n* (FOV = 1.00 – 0.85 mm when *n* = 1.33 – 1.56), which is not clipped by the 1.00 mm FOV of the 20X objective (O3). The physical layout, ZEMAX model, and objective options for this FOV-maximized NODO imaging configuration are shown in **Supplementary Figure 11**. See **Supplementary Note 2** for more discussion of NA-versus FOV-maximized imaging. As mentioned previously, an alternative design could use the CFI S Plan Fluor LWD 20XC and a customized O3 to make use of the full 0.483 NA of our O1 (see **Supplementary Figure 12** and **Supplementary Note 5**).

### ODO optical path

The physical layouts and ZEMAX models of the ODO imaging configurations are shown in **Supplementary Figure 13-14**. While the NODO imaging path can provide sub-micrometer resolution, the FOV of the multi-immersion primary objective (O1) is restricted to 1 mm, which is insufficient for fast mesoscopic imaging. This tradeoff between NA and FOV is standard across all currently available microscope objectives (e.g., no current clearing-compatible objectives can simultaneously offer sub-micrometer resolution over a mesoscopic FOV) (**Supplementary Figure 8**) [49]. Therefore, in our hybrid system, we achieve low-resolution imaging with a second independent ODO imaging path.

The ODO collection path uses the same objective as the illumination optical path (TL2X-SAP, Thorlabs). The objective is similarly used in conjunction with a SIMlens (fabricated by BMV Optical), which provides multi-immersion performance and prevents axial chromatic aberrations in the ODO imaging path [26]. In addition, the SIMlens increases the NA of the collection path by a factor of *n*. This yields an effective NA = 0.10 × *n*.

The current set point of the ODO collection path lies between the NA- and FOV-maximized imaging extremes and uses a tube lens with a 200-mm focal length (TTL200-A, Thorlabs). This provides a magnification of 2 × *n* and an effective FOV of 6.6 mm / *n*, which undersamples the collection NA of the system when using a sCMOS camera with a pixel spacing of 6.5 μm (pco.edge 4.1, PCO Tech, 2048 × 2048 pixels). In this configuration, the back aperture of the illumination objective is underfilled to yield an illumination NA of ∼0.04 × *n*. This results in a confocal parameter of ∼250 – 300 μm, which corresponds to a digital image height of ∼128 pixels for each raw camera image. The width (in pixels) of each raw camera image is the full 2048 pixels of the sCMOS camera, corresponding to a distance of ∼4.25 – 5 mm in the sample. This combination of illumination and collection NA provides near-isotropic resolution (**Supplementary Figure 16**Error! Reference source not found.).

The magnification of the ODO imaging path is easily adjusted by changing the tube lens. NA-maximized imaging is achieved using a tube lens with a 400-mm focal length (AC508-400-A, Thorlabs). This provides a magnification of 4 × *n* and an effective FOV of 3.3 mm / *n*, which corresponds to a near-Nyquist sampling rate of ∼2.1 when using a sCMOS camera with a pixel spacing of 6.5 μm. FOV-maximized imaging is achieved using a tube lens with a 100-mm focal length (TTL100-A, Thorlabs). This provides a magnification of 1 × *n* and an effective FOV of 13.33 mm / *n*. See **Supplementary Note 2** for more discussion of NA- versus FOV-maximized imaging modes.

### Image acquisition and post processing

Image strips are collected with a combination of stage-scanning and lateral/vertical tiling. The stage-scanning firmware is used to send a TTL SYNC trigger signal from the XY stage to the sCMOS camera for reproducible initiation of each imaging strip. After triggering, the camera is set to free-running mode and acquires the desired number of frames for a given image-strip length (as the sample is scanned by the stage at a constant velocity), with a spacing between adjacent frames that is identical to the sampling pitch of the raw camera images. For each raw frame, the camera uses the standard rolling shutter, where the shutter rolls from the center of the camera chip in both directions, as opposed to the light-sheet readout mode, where the shutter rolls from the top to the bottom of the camera chip. Using the standard rolling shutter, the rolling directions are oriented along the light sheet propagation direction. This orientation allows the imaging speed to be increased when the pixel height of each raw frame is cropped to match the confocal parameter of the illumination light sheet.

At the start of each exposure, the camera sends a trigger to an analog output DAQ card (PCIe-6738, National Instruments). The DAQ card then sends output voltages to the lasers, galvos, and ETL (for alignment only, not axial sweeping). To reduce motion blur, the lasers and galvos are triggered with a delay to only illuminate once the shutter has rolled across the full pixel height of the camera frame, resulting in a strobing effect. For the 128- or 256-pixel height of the ODO and NODO paths respectively, the rolling time, *t*_roll_, is 0.625 and 1.25 ms. The exposure time, *t*_exp_, is set to 3 × *t*_roll_, resulting in a total exposure time, t_tot_, of 4 × *t*_roll,_ or 2.5 and 5 ms for the ODO and NODO paths. This corresponds to a data rate that is 1/4 the maximum data rate of the sCMOS camera. The lateral scanning mirror is actuated with a sawtooth waveform and completes a single period within the total exposure time, *t*_tot_, of the raw camera frame, corresponding to a frequency of ∼400 and ∼200 Hz for the ODO and NODO paths. The ETL and second mirror are set to a pre-calibrated DC voltage for the entire image strip to yield an in-focus light sheet that is axially aligned to the center of the camera chip. Raw camera frames are streamed from the camera to RAM and subsequently saved directly as a single HDF5 file and XML file with the associated metadata, which is suitable for immediate processing with BigStitcher via ImageJ [50, 51]. This involves on-the-fly saving of down-sampled copies of the image strip (2x, 4x, 8x, and 16x) in the hierarchical HDF5 file format. Additionally, the GPU-based B3D compression filter can be optionally added to the HDF5 writing process to yield 5-10X compression with negligible loss of usable information content [52]. However, both processing steps slow down the net data rate from RAM to disk. On-the-fly saving of down-sampled copies of the image strips slows the data rate by ∼2x, and the B3D algorithm slows down the process by another ∼2x. This net speed reduction of ∼4x motivated our selection of the camera frame rate mentioned previously. For our experiments, this reduction in post-processing times and data-storage requirements are worth the reduction in imaging speed. However, it is important to note that the imaging speeds could be increased by 2x if the aforementioned processing steps are omitted from the acquisition procedure, in which case the imaging speed would become limited by the duty cycle of the illumination strobing, where *t*_exp_ = *t*_*roll*_. The imaging speeds quoted in **Figure 1g** assume the 4x-reduced data rate.

A ∼15% overlap is used for both vertical and lateral tiling. Different wavelength channels are acquired sequentially. For each image strip “tile” that is acquired by laterally scanning the specimen in the *y*-direction, all channels are acquired by cycling through various laser/filter combinations and re-scanning that image strip for each laser/filter setting, before moving to the next tile position. When tiling vertically, the laser power is increased with depth per a user-defined exponential relationship, *P* = *P*_0_ × exp(*z*/*μ*), to account for the attenuation of the illumination light sheet as it penetrates deeper into the specimen (typically *μ* = 5 – 20 mm^-1^). Finally, if desired, all of the imaging tiles can be aligned and fused into one contiguous 3D image as an HDF5 or TIFF file output using BigStitcher [50]. The entire image acquisition is controlled using a custom-written Python program that is available from the authors upon request.

### Computer hardware

During acquisition, the images are collected by a dedicated custom workstation (Puget Systems) equipped with a high-specification motherboard (Asus WS C422 SAGE/10G), processor (Intel Xeon W-2145 3.7GHz 8 Core 11MB 140W), and 256 GB of RAM. The motherboard houses several PCIe cards, including 2 CameraLink frame grabbers (mEIV AD4/VD4, Silicon Software) for streaming images from the camera, a DAQ card (PCIe-6738, National Instruments) for generating analog output voltages, a 10G SFP+ network card (StarTech), and a GPU (TitanXP, NVIDIA). Datasets are streamed to a local 8 TB U.2 drive (Micron) that is capable of outpacing the data rates of the microscope system. Data is then transferred to a mapped network drive located on an in-lab server (X11-DPG-QT, SuperMicro) running 64-bit Windows Server, equipped with 768 GB RAM and TitanXP (NVIDIA) and Quadro P6000 (NVIDIA) GPUs. The mapped network drive is a direct-attached RAID6 storage array with 15 × 8.0 TB HDDs. The RAID array is hardware based and controlled by an external 8-port controller (LSI MegaRaid 9380–8e 1 GB cache). Both the server and acquisition workstation are set with jumbo frames (Ethernet frame), and parallel send/receive processes matched to the number of computing cores on the workstation (8 physical cores) and server (16 physical cores), which reliably enables ∼ 1.0 GB sec^−1^ network-transfer speeds.

### Preparation of ECi-cleared mouse brain

Labeling and clearing was carried out as previously described [43]. For sparse labeling, a Slc17a7-Cre mouse (8 weeks old, female) received a systemic injection, via the retro-orbital sinus, of a mixture of Cre-dependent Tet transactivator (PHP-eB-Syn-Flex-TRE-2x-tTA) and a reporter virus (PHP-eB-CAG-TRE-3xGFP) [43, 53]. High-titer (> 10^12^ GC/mL) viruses were obtained from the Janelia Research Campus Molecular Biology Core and diluted in sterile water when necessary.

Transfected mice were anesthetized with an overdose of isoflurane and then transcardially perfused with a solution of PBS containing 20 µg/mL heparin (Sigma-Aldrich #H3393) followed by a 4% paraformaldehyde solution in PBS. Brains were extracted and post-fixed in 4% paraformaldehyde at 4°C overnight (12-14 hrs) and washed in PBS to remove all traces of excess fixative (PBS changes were performed at 1 hr, 6 hr, 12 hr, and 1 day).

For amplification by immuno-labeling, brains were delipidated with a modified Adipo-Clear protocol [54]. Brains were washed with a methanol gradient series (20%, 40%, 60%, 80%, Fisher #A412SK) in B1n buffer (H_2_O/0.1% Triton X-100/0.3 M glycine, pH 7; 4 mL / brain; one hr / step). Brains were then immersed in 100% methanol for 1 hr, 100% dichloromethane (Sigma #270997) for 1.5 hrs, and three times in 100% methanol for 1 hr. Samples were then treated with a reverse methanol gradient series (80%, 60%, 40%, 20%) in B1n buffer for 30 min each. All procedures were performed on ice. Samples were washed in B1n buffer for one hr and left overnight at room temperature; and then again washed in PTxwH buffer (PBS/0.1% Triton X-100/0.05% Tween 20/2 µg/mL heparin) with fresh solution after one and two hrs and then left overnight.

After delipidation, selected samples were incubated in primary antibody dilutions in PTxwH for 14 days on a shaker (1:1000, anti-GFP, Abcam, #ab290). Samples were sequentially washed in 25 mL PTxwH for 1, 2, 4, 8, and three times for 24 hrs. Samples were incubated in secondary antibody dilutions in PTxwH for 14 days (1:600, AlexaFluor® 488 conjugated donkey-anti-rabbit IgG) and washed in PTxwH similar to descriptions above. Finally, the tissue was dehydrated in ethanol grades (25, 50, 75, 100%) for 8 hr per grade. The 100% grade was repeated to ensure removal of all water from the tissue. Finally, the tissue was cleared in ethyl-cinnamate (Sigma-Aldrich #112372) for 8 hr before imaging. All experimental protocols were conducted according to the National Institutes of Health guidelines for animal research and were approved by the Institutional Animal Care and Use Committee at Howard Hughes Medical Institute, Janelia Research Campus.

### Preparation of MDA-231/OS-RC-2 and αSMA-labeled mouse brains

Human breast cancer cells, MDA-MB-231-5a-D (MDA-231) are highly metastatic clones from MDA-MB-231 [55]. Human renal cell carcinoma, OS-RC-2 were kindly provided by Prof. Tatsuro Irimura (Juntendo University, Japan). MDA-231 cells were cultured in Dulbecco’s modified eagle’s medium (DMEM) supplemented with 10% fetal bovine serum (FBS), 100 U/ml penicillin and 100 µg/ml streptomycin as previously described [55]. OS-RC-2 cells were maintained in RPMI1640 containing 10% FBS and penicillin/streptomycin [56].

To establish cancer cells stably expressing firefly luciferase and mCherry under the EF-1 promoter, a lentiviral expression system was used (kindly provided by Dr Hiroyuki Miyoshi, deceased, formerly Keio University, Tokyo, Japan) as described previously (Miyoshi et al., Proc. Natl. Acad. Sci., 1997). Briefly, 293FT cells were transfected with a vector construct encoding the expression protein, VSV-G, a Rev-expressing construct (pCMV-VSV-G-RSV-Rev), and a packaging construct (pCAG-HIVgp). The culture supernatants containing viral particles were collected and used as lentiviral vectors.

BALB/c-nu/nu mice (4 weeks old, female) were purchased from Japan SLC (Shizuoka, Japan). All experiments were approved and carried out according to the Animal Care and the Use Committee of the Graduate School of Medicine, The University of Tokyo. For developing experimental brain metastasis models by intracardiac (i.c.) inoculation, BALB/c-nu/nu mice were injected with MDA-231 or OS-RC-2 cells (5 × 10^5^ cells /mouse) by puncture into the left ventricle of the heart.

Clearing, 3D staining, and imaging of whole mouse brain samples was performed with CUBIC clearing and CUBIC-HistoVision (CUBIC-HV) staining protocols [7, 10]. An updated CUBIC-HV staining protocol was used (HV1.1, commercialized by CUBICStars Co. and Tokyo Chemical Industry (TCI), TCI #C3709, #C3708). In brief, the PFA-fixed whole mouse brains were treated with CUBIC-L for 4 days at 37 °C, washed with PBS, stained with SYTOX-G (1/2500) in CUBIC-HV nuclear-staining buffer (included in TCI #C3709) for 5 days at 37 °C, and then washed / immersed in 50 % and 100% CUBIC-R+ for 1 day and 3 days, respectively, at room temperature. We used CUBIC-R+(M) [45 wt% of antipyrine (TCI #D1876), 30 wt% of N-methylnicotinamide (TCI #M0374), and 0.5% (v/w) N-butyldiethanolamine (TCI #B0725), adjusted to pH ∼10] for brains containing cancer cells, and CUBIC-R+(N) [45 wt% of antipyrine, 30 wt% of nicotinamide (TCI #N0078), and 0.5% (v/w) N-butyldiethanolamine, adjusted to pH ∼10] for the αSMA immunostained brain, respectively.

For the whole mouse brain immunostained with anti-α-SMA (Sigma, #A5228) antibodies, the brain was subjected to CUBIC-HV immunostaining. The brain was first treated with 3 mg/mL hyaluronidase in CAPSO buffer (pH 10) for 24 h at 37°C. After washing with hyaluronidase wash buffer [50 mM Carbonate buffer, 0.1% (v/v) Triton X-100, 5% (v/v) Methanol, and 0.05% NaN3] and HEPES-TSC buffer [10 mM HEPES buffer, pH 7.5, 10% (v/v) Triton X-100, 200 mM NaCl, 0.5% (w/v) casein, and 0.05% NaN3], the brain was immersed in 500 µL of HEPES-TSC buffer containing a primary antibody (6 µg for anti-α-SMA), a secondary Fab fragment (FabuLight, Jackson immunolab, Alexa Fluor® 594 Goat Anti-Mouse IgG1 #115-587-185, Alexa Fluor® 594 Goat Anti-Rabbit IgG #111-587-008, Alexa Fluor® 594 Goat Anti-Mouse IgG2a #115-587-186, 1:0.75 of weight ratio), and 3D immunostaining additive (1x) (included in TCI #C3708). Then, the sample was incubated with gentle shaking for 10 days at 32 °C. After staining, the sample was additionally incubated in the same buffer for 1 day at 4°C to stabilize the Fab binding. Then, the sample was washed and post-fixed according to the protocol of CUBIC-HV immunostaining kit (TCI #C3708) before being index-matched with CUBIC-R+. The cleared sample was embedded in CUBIC-R-agarose for imaging and storage [57]. This animal experimental procedures and housing conditions of the animals were approved by the Animal Care and Use Committees of the Graduate School of Medicine of the University of Tokyo.

### Statistics and reproducibility

For statistical analysis, we reported the mean, standard deviation, and number of observations. For repeatability, only one sample was used for each imaging experiment, unless otherwise noted.

## Supporting information

ZEMAX Files

Objective Lens Data

Video Abstract

## Data Availability

The customized ZEMAX files are available as **Supplementary Data**. Imaging datasets are available from the authors upon request.

## Code Availability

The simulation codes used to model the lateral and axial resolution of the various microscope architectures is available on [GitHub](https://github.com/adamkglaser/OPM_simulations) and as **Supplementary Code**. The acquisition software code for the microscope is available from the authors upon request.

## Acknowledgements

We would like to thank Andrew York and Alfred Millet-Sikking for discussions regarding oblique planar microscopy, remote focus imaging, and alignment procedures. We would also like to thank Jon Daniels for discussions and his development of the multi-immersion objective. We would also like to thank Chika Shimizu (RIKEN BDR) for the support of preparing CUBIC-cleared and stained specimens and Kohei Miyazono (The University of Tokyo) for the discussion and support of brain metastasis experiments. This work was funded in part by the National Institutes of Health (NIH) K99 CA240681 (Glaser), R01CA244170 (Liu), R01EB031002 (Liu), R01GM079712 (Imaizumi), R01DK107436 (Xin), R01DK092202 (Xin); Department of Defense (DoD) Prostate Cancer Research Program (PCRP) W81XWH-18-10358 (Liu and True), W81XWH-19-1-0589 (Reder), W81XWH-20-1-0039 (Wei), and Prostate Cancer Young Investigator Award (Reder); National Science Foundation (NSF) Graduate Research Fellowship DGE-1762114 (Barner and Bishop); NSF 1934292 HDR: I-DIRSE-FW (Liu and Serafin); Washington Research Foundation Postdoctoral Fellowship (Stoltzfus); Science and Technology Platform Program for Advanced Biological Medicine by the Japan Agency for Medical Research and Development (AMED) JP21am0401011 (Ueda), ERATO by Japan Science and Technology Agency (JST) JPMJER2001 (Ueda), HFSP Research Grant Program RGP0019/2018 (Ueda), AMED-PRIME JP21gm6210027 (Susaki), Grants-in-Aid for Scientific Research on Innovative Areas (JSPS KAKENHI grant) 17H06328 (Susaki), Grants-in-Aid for Scientific Research on Innovative Areas (JSPS KAKENHI grant) 20K1612 (Kubota). Work in the Murawala laboratory is supported by grants from NIH-COBRE (5P20GM104318-08) and DFG (429469366).

## Author Contributions

A.K.G. and J.T.C.L. conceived of and designed the microscope system. P.R.N. provided feedback on the system design and its potential applications. A.K.G., K.W.B., R.B.S., and G.G. performed simulations of the microscope. A.K.G. fabricated the microscope system with help from L.A.B. E.A.S. and H.R.U. provided and prepared the immunostained CUBIC-cleared mouse brains. E.A.S., S.I.K., and H.R.U. prepared and provided the metastatic mouse brains. J.C. and K.S. prepared and provided the mouse brain. P.B., E.T., and R.C.R. provided the mouse brain with preparation by A.K.G. The human brain slice was prepared by A.K.G. H.L. and L.A.G.L. provided quantitative analysis of the high-resolution images. Y.Y. and H.Z. provided and prepared the PEGASOS-cleared mouse brain. E.K.N., B.J.B., and J.S. provided and prepared the SHIELD-cleared mouse embryos. H.H., N.P.R., and L.D.T. provided and prepared the ECi-cleared human prostate tissue. J.J.W., R.S., E.S., C.R.S., and M.Y.G. provided and prepared the Ce3D-cleared mouse lymph node. X.W. and L.X. provided and prepared the iDISCO-cleared mouse prostate. A.K.H. and T.I. provided and prepared the ClearSee-processed Arabidopsis plant and M.P., P.M., and H.U.D. provided and prepared the DEEP-Clear-processed Axolotl. A.K.G. and J.T.C.L. led the writing of the manuscript. All authors contributed to the manuscript.

## Corresponding Author

Correspondence to <u>Adam K. Glaser</u> or <u>Jonathan T.C. Liu</u>

