## Supplementary Material for "A hybrid open-top light-sheet microscope for multi-scale imaging of cleared tissues"

*Supplementary Note 1 – Hybrid system design choices*

We previously developed a number of open-top light-sheet (OTLS) microscopy systems that provide an ease-of-use similar to a flat-bed document scanner, enabling one or more specimens of arbitrary geometry to be imaged without lateral constraints at depths of up to several millimeters [1-4]. The OTLS architecture also accommodates a wide range of potential accessory technologies, such as microfluidics, electrophysiology, and micro-dissection/aspiration. However, in addition to being somewhat limited in imaging depth (e.g., 2-3-mm deep for a recent system) [4], these systems have had strict index-matching requirements between the specimen and specimen holder when operating at the higher numerical apertures (NA) needed to achieve moderate-resolution imaging (**Supplementary Figure 1**) [4]. Furthermore, these OTLS prototypes have largely been constrained to image at fixed-resolution set points (low to moderate) [1, 2]. Since volumetric imaging speeds scale to the third power with respect to spatial resolution, we were motivated by the need for a system that could enable a hierarchical multi-scale imaging workflow, as is common with 2D microscopes containing a turret of objectives. While multi-resolution 3D microscopy systems have been developed in the past [4], they have not featured the resolution range, imaging depth, and tolerance to index mismatch (ease of use) that we have desired. Therefore, we sought to design a new OTLS microscope system that could serve as a versatile multi-scale imaging platform for a wider array of cleared-tissue imaging applications.

Our prior generations of OTLS microscopes featured an orthogonal dual objective (ODO) architecture (similar to inverted light-sheet systems), in which the illumination and collection paths are each oriented at 45 deg with respect to the vertical (gravitational) axis [1-8]. While these systems achieved varying levels of resolution (several µm to sub-micrometer), there were two major design trade-offs with the ODO architecture as we sought to improve spatial resolution. (1) High-NA objectives tend to have shorter working distances that limit imaging depth in thick specimens, which is exacerbated if the objectives are oriented at an oblique angle with respect to a specimen holder [4]. (2) In the case of open-top systems at high NAs, the quality of off-axis focused beams is severely degraded (aberrated) in the presence of minute refractive-index disparities between the cleared tissue, sample holder, and immersion medium [2, 4, 5].

To overcome these design trade-offs, we considered the use of a single-objective light-sheet architecture that has gained popularity in recent years [9-21]. With single-objective light-sheet systems, the illumination and collection beams share a single objective, in which a remote focus is used to re-image the non-orthogonal light sheet onto a flat detector array (camera chip). Orienting the objective in the normal direction (perpendicular) with respect to the specimen holder makes full use of the objective working distance and dramatically reduces the system's sensitivity to refractive-index disparities. While the single-objective light-sheet architecture mitigates the working-distance and index-mismatch limitations of ODO, there are several shortcomings of single-objective light-sheet microscopy. Because the illumination and collection paths share one objective, their NAs and crossing angle are limited and are also coupled to one another such that axial and lateral resolution must trade off (**Supplementary Figure 3**) [22]. While this is less problematic for objectives with very high NAs (e.g., >1.0), such objectives exhibit a limited field of view (FOV) and short working distance, which are not ideal for multi-scale imaging of large cleared tissues (**Supplementary Figure 8**) [23]. One advantage of single-objective light-sheet systems is that they facilitate the use of a single mirror to scan and de-scan the illumination and collection paths for imaging live, dynamically changing specimens. However, this ability is less important for imaging fixed tissues, in which stage scanning is a viable alternative.

To achieve an optimal combination of resolution (axial and lateral), working distance, and insensitivity to index-mismatch, we developed the NODO architecture [22]. By using a separate objective for non-orthogonal illumination, which allows the full NA of a vertically oriented objective to be used for fluorescence collection (with the same remote-focus imaging strategy as single-objective light-sheet microscopy), our NODO configuration maintains the working-distance and index-mismatch-insensitivity advantages of single-objective light-sheet microscopy, while offering the improved imaging performance of a dual-objective light-sheet system (**Supplementary Figures 3-4**). The use of a separate illumination objective increases the crossing angle of the two beams, which reduces the angle of the remote objective that is used for re-imaging the tilted light sheet. This enables standard off-the-shelf air objectives to be used as a remote objective rather than specialized bespoke objective assemblies (**Supplementary Figure 7**) [13, 14, 17, 18]. Finally, our selection of a vertically oriented primary objective with optimal optical and mechanical specifications allows for the incorporation of a separate collection arm at 45 deg for low-resolution OTLS imaging via an ODO architecture (using the same illumination optics as the NODO path). In this ODO imaging path, a low-NA collection objective with a large FOV is used to enable rapid mesoscopic-resolution imaging.

*Supplementary Note 2 – NA- and FOV-maximized imaging*

Both collection objectives in the NODO and ODO paths provide more imaging information (i.e., space-bandwidth product) than current 2048 x 2048 pixel sCMOS cameras can capture. Therefore, the multi-scale capabilities of our system can be extended by changing the magnification (i.e., FOV) of both imaging paths on the sCMOS chip; either by zooming in until the imaging resolution is limited by the NA of the objective (i.e. "NA-maximized" imaging), or by zooming out until the tissue area that is imaged on the sCMOS camera chip is limited by the FOV of the objective (i.e. "FOV-maximized" imaging). In the latter case, if the effective NA of the objective is not reduced, the resolution is determined by the pixel spacing (i.e., sampling ratio) on the sCMOS camera, according to the Nyquist criterion. However, in practice, imaging with a higher NA than necessary can introduce aberrations at the edges of the larger FOV. This is especially true for the ODO imaging path, where the SIMlens provides diminishing performance farther from the center of the FOV [3]. Therefore, for both the NODO and ODO paths, we reduce the effective NA to provide near-Nyquist sampling over the increased FOV. For the NODO path, this is achieved by switching O3 at the remote focus (**Supplementary Figure 7**), and for the ODO path this is achieved using a variable aperture near the back focal plane behind the collection objective (**Supplementary Figures 13-14**).

*Supplementary Note 3 – Comparison to LSTM*

Note that the physical layout of our hybrid system bears some resemblance to the light-sheet theta microscopy (LSTM) architecture, which employs dual-path illumination with a pair of off-axis objectives [24]. However, the NODO architecture, which forms one part of our multi-scale hybrid system, is >10X more light-efficient compared to LSTM with fewer trade-offs between light efficiency and axial resolution

(**Supplementary Figure 31**). Instead of using a remote focus to re-image a tilted light sheet, LSTM scans both beams (with a complex and synchronized combination of lateral and axial scanning) to generate a virtual light sheet within the specimen that is orthogonal to the collection objective. To reject out-of-focus light, the scanning beams are synchronized to a confocal slit, which results in inefficient light collection. This is in contrast to other light-sheet microscopy architectures where the imaging plane is always co-planar with the light sheet.

*Supplementary Note 4 – Multi-scale imaging with the single-objective light-sheet architecture*

With currently available commercial microscope objectives (**Supplementary Figure 8**), multi-scale imaging at both mesoscopic and sub-micrometer resolution with a single-objective light-sheet microscopy architecture is not possible. This is because no single objective currently provides sub-micrometer resolution over a mesoscopic FOV. Furthermore, it is not feasible to simply add a lower-magnification imaging objective to an existing single-objective light-sheet system (i.e., a NODO system in which the light sheet is still delivered through the primary objective and a second non-orthogonal imaging objective is used as a second imaging path). This is because the light sheet width would be constrained by the FOV of the primary objective and would not be able to take advantage of the larger FOV of the lower-magnification objective that is used to create the remote focus, O2. Therefore, multi-scale imaging with the single-objective light-sheet architecture may only be possible in the future with the development of highly customized objective (see **Supplementary Figure 29**). While these objectives would likely be prohibitively expensive to design and manufacture, the patent literature does contain recent examples of highly customized objectives designed specifically for whole slide scanners (e.g., US #8350904 which discloses a 0.90 NA objective with a 1.5 mm FOV and ~1 mm working distance [23]). However, the working distance is low and this would require the relay lenses and both remote objectives to accommodate the same large FOV and high NA, which is not ideal for multi-scale imaging.

*Supplementary Note 5 – Alternative NODO architecture*

An alternative NODO configuration could use the CFI S Plan Fluor LWD 20XC (Nikon) and a customized O3 to maintain the full 0.483 NA of our O1. Several custom options for O3 include 20X water-immersion objectives from Olympus and Zeiss, as well as the recently developed solid immersion AMS-AGY v1.0 objective from Special Optics. While the AMS-AGY v1.0 (Special Optics) is a convenient option, the FOV of the objective is limited to 0.15 mm (diffraction-limited) and 0.25 mm (near diffraction-limited), which is too small for our design. On the other hand, using a water-immersion objective requires sealing the water into a custom housing or prism with a cover glass. This is not compatible with the Olympus water objective, as it is not designed for use with a cover glass. Therefore, the only viable option is the Zeiss "W" Plan Apochromat 20X. We developed but did not ultimately configure and test a custom sleeve to house this objective, which seals water around the tip of the objective using a gasket and cover glass. When used for O3 and combined with a downstream zoom module (for example the Resolv4K, Navitar), this configuration could simultaneously enable both NA- and FOV-maximized imaging. While we will continue to explore and test this configuration, these components add cost and complexity to the design. The physical layout, ZEMAX model, and the customized Zeiss water objective assembly are shown in **Supplementary Figure 12**. Note, an updated AMS-AGY v2.0 objective is now available with a larger 0.45 mm field of view [18].

*Specimen holders*

Transparent specimen-holder materials can be interchanged and fixed to the bottom of a modular holder to ensure adequate index matching for different clearing protocols (FEP or PTFE for expanded gels or ClearSee, fused silica for CLARITY, PMMA for Ce3D, BK7 glass for CUBIC, COC or COP for PEGASOS, and HIVEX for iDISCO, BABB, or ECi) [2]. See **Supplementary Figure 5** for a full matrix of specimen holder materials and clearing protocols.

*Optical performance simulations*

To calculate the axial and lateral resolution of the various system configurations, a custom simulation code was developed in Python. In short, the polar and azimuthal ray angles (spherical coordinate system) for a given NA across the pupil of O2 are calculated over a grid with a user-defined numerical resolution (default 256 x 256). The incident angle of these rays on a tilted O3 are then calculated, as shown in **Supplementary Figure 30** and described in [5]. Based on the acceptance NA of O3, the pupil plane is then clipped in 2D, yielding the final pupil pattern $P(k_x,k_y)$. Finally, the 3D point spread function (PSF) can be calculated as:

$$PSF(x, y, z) = \iint P(k_x, k_y)\, e^{2\pi i(k_x x + k_y y)} e^{2\pi i k_z(k_x,k_y)z} d_{k_x} d_{k_y}$$

where $k_z(k_x,k_y)$ acts to refocus the pupil pattern at varying axial positions [25]. Once the 3D PSFs are calculated, the full-width-half-maximum (FWHM) dimensions of the PSF can be obtained along the three orthogonal directions (xyz). This code is available as **Supplementary Code**.

*Index-mismatch simulations*

To quantify the index-mismatch sensitivity of the various system configurations, simulations were performed using the optical ray-tracing software OpticStudio (ZEMAX). Blackbox files for the multi-immersion objective (#54-12-8, Special Optics) and 2X air objective (TL2X-SAP, Thorlabs) were used. The refractive index of the immersion medium and specimen were set to $n$ = 1.46. The optical path length of the specimen holder ($\Delta n$ x $t$) was then varied. To assess the PSF, the real and imaginary components of the electric field were exported from ZEMAX, and input into a MATLAB (Mathworks) implementation of the beam propagation method (BPM) to obtain a full 3D PSF [26]. The Strehl ratio was evaluated by finding the maximum value of the 3D PSF.

*Preparation of Chat-IRES-Cre;Ai162 cleared brain*

An adult (p100) Chat-IRES-Cre;Ai162 mouse was transcardially perfused with 4% paraformaldehyde (PFA); the brain was extracted and postfixed in 4% PFA for four hours prior to short-term storage in 1M PBS. The brain was then CUBIC cleared using the same protocol as the MDA-231/OS-RC-2 brains with CUBIC-R+(N) to preserve GCaMP6s fluorescence. All procedures involving transgenic mice were conducted in accordance with NIH guidelines and approved by the Institutional Animal Care and Use Committee (IACUC) of the Allen Institute for Brain Science.

*Preparation of CUBIC-cleared human brain slab*

Human brain tissue from an Alzheimer's disease donor was obtained from the University of Washington Biorepository and Integrated Neuropathology (BRaIN) Laboratory. The brain tissue was cleared using the CUBIC clearing protocol [27]. In brief, the fixed human brain tissue was treated with 500 mL of CUBIC-L for 4 weeks at 37 °C, refreshing the CUBIC-L solution every 3-4 days. The tissue was then washed with PBS and stained with 30 µM pFTAA (Millipore #SCT066) in 200 mL of Sca/eCUBIC-1A [10% w/v Triton-X100 (Sigma #X100-500ML), 5% w/v Quadrol (Sigma #122262-1L), 10% w/v urea (Sigma # 15604-250G)] with 500 mM NaCl (Fisher #18-606-422) for 2 weeks a\t 37 °C. After staining, the tissue was washed / immersed in 50 % and 100% CUBIC-R+(N) [45 wt% of antipyrine (Fisher #AC104975000), 30 wt% of nicotinamide (Fisher # N0078500G), and 0.5% (v/w) N-butyldiethanolamine (Sigma # 471240-500ML), adjusted to pH ~10] for 3 day and 5 days, respectively, at room temperature. The cleared brain tissue was finally embedded in CUBIC-R-agarose for imaging and storage [28].

*Preparation of PEGASOS-cleared mouse brain*

Thy1-EGFP mouse brain samples were processed with the PEGASOS tissue-clearing method following an established protocol [29]. Briefly, mice were anesthetized with an intraperitoneal injection of a combination of xylazine and ketamine anesthetics (Xylazine 10-12.5 mg/kg; Ketamine, 80–100 mg/kg body weight), and transcardially perfused with 50 ml ice-cold heparin PBS to wash off the blood, and then with 50 ml 4% PFA for fixation. Mouse brains were dissected and post-fixed in 4% PFA at room temperature for 24h. Subsequently, tissues were decolorized with decolorization solution [25% (v/v in $H_2O$) Quadrol (Sigma-Aldrich #122262)] in a 37°C shaker for two days. Then delipidation was performed in a 37°C shaker using a tert-Butanol gradient at 6-12 hours for each step: 30% tert-Butanol (tB, Sigma-Aldrich #471712) solution (70% v/v $H_2O$, 27% v/v tB, 3% w/v Quadrol), 50% tB solution

(50% v/v $H_2O$, 47% v/v tB, 3% w/v Quadrol), and 70% tB solution (30% v/v $H_2O$, 67% v/v tB, 3% w/v Quadrol). Following that, mouse brains were immersed into a dehydration solution (70% tB, 27% (v/v) poly(ethylene glycol) methyl ether methacrylate average Mn500 (PEG MMA500) (Sigma-Aldrich #447943) and 3% (w/v) Quadrol) in a 37°C shaker for 2 days. The final clearing medium was composed of 75% (v/v) benzyl benzoate (BB) (Sigma-Aldrich #B6630), 22% (v/v) PEG MMA500 and 3% (w/v) Quadrol. After dehydration, mouse brains were incubated in this clearing medium in a 37°C shaker until they turned fully transparent. Cleared samples were preserved in clearing medium at room temperature for storage and imaging. Procedures involving mice were approved by the Institutional Animal Care and Use Committee of Texas A&M in accordance with NIH guidelines.

*Preparation of CUBIC-cleared immunolabeled mouse brains*

Clearing, 3D staining and imaging of whole mouse brain samples were performed with CUBIC clearing and CUBIC-HistoVIsion (CUBIC-HV) staining protocols [27, 30]. We applied an updated CUBIC-HV staining protocol (HV1.1, commercialized by CUBICStars Co. and Tokyo Chemical Industry (TCI), TCI #C3709, #C3708). In brief, the PFA-fixed whole mouse brains were treated with CUBIC-L for 4 days at 37 °C, washed with PBS, stained with SYTOX-G (1/2500) in CUBIC-HV nuclear-staining buffer (included in TCI #C3709) for 5 days at 37 °C, and washed / immersed in 50 % and 100% CUBIC-R+(N) [45 wt% of antipyrine, 30 wt% of nicotinamide (TCI #N0078), and 0.5% (v/w) N-butyldiethanolamine, adjusted to pH ~10]for 1 day and 3 days, respectively, at room temperature.

The whole mouse brains immunostained with anti-PV (Swant, #PV235) and anti-ChAT (Abcam, #ab178850) antibodies were subjected to CUBIC-HV immunostaining. The nuclear-labeled sample was treated with 3 mg/mL hyaluronidase in CAPSO buffer (pH 10) for 24 h at 37°C. After washing with hyaluronidase wash buffer [50 mM Carbonate buffer, 0.1% (v/v) Triton X-100, 5% (v/v) Methanol, and 0.05% $NaN_3$] and HEPES-TSC buffer [10 mM HEPES buffer, pH 7.5, 10% (v/v) Triton X-100, 200 mM NaCl, 0.5% (w/v) casein, and 0.05% $NaN_3$], the brain was immersed in 500 µL of HEPES-TSC buffer containing a primary antibody (10 µL for anti-PV, 5.3 µg for anti-ChAT), a secondary Fab fragment (FabuLight, Jackson immunolab, Alexa Fluor® 594 Goat Anti-Mouse IgG1 #115-587-185, Alexa Fluor®

594 Goat Anti-Rabbit IgG #111-587-008, Alexa Fluor® 594 Goat Anti-Mouse IgG2a #115-587-186, 1:0.75 of weight ratio), and a 3D immunostaining additive (1x) (included in TCI #C3708, for the ChAT staining). Then, the sample was incubated with gentle shaking for 17 days (anti-PV) at 32 °C or 10 days (anti-ChAT) at room temp. After staining, the sample was additionally incubated in the same buffer for 1 day at 4°C to stabilize the Fab binding. Then, the sample was washed and post-fixed according to the protocol of CUBIC-HV immunostaining kit (TCI #C3708) before the RI-matching by CUBIC-R+. The cleared sample was embedded in CUBIC-R-agarose for imaging and storage. The cleared sample was embedded in CUBIC-R-agarose for imaging and storage [28]. This animal experimental procedures and housing conditions of the animals were approved by the Animal Care and Use Committees of the Graduate School of Medicine of the University of Tokyo.

*Preparation of DEEP-Clear-processed axolotl*

Axolotl samples were first anesthetized with benzocaine, followed by fixation with 4% PFA at 4°C overnight. Specimens were washed several times with PBS at room temperature to remove fixative, followed by pre-chilled acetone treatment at -20°C overnight. After three PBS washes at room temperature (gentle shaking), specimens were incubated in Solution-1 (axolotl) at 37°C for three hours under gentle shaking. Multiple PBS washes were used to remove Solution-1 from tissues, and samples were mounted in Solution-2 for light-sheet imaging.

Solution-1 consists of 10% (v/v) THEED (Sigma-Aldrich #87600-100ML), 5% (v/v) Triton® X 100 (Roth #3051.2) and 25% (w/v) Urea (Roth #X999.2) in d$H_2O$ e.g., 5 mL THEED, 2.5 mL Triton® X 100 and 12.5 g Urea for a 50 mL solution volume.

Solution-2 consists of a 50% (w/w) meglumine diatrizoate (Sigma-Aldrich #M5266) in PBS (pH 9-9.3) (e.g., 10 g of meglumine diatrizoate added to 10 mL of PBS). The final refractive index was further adjusted to 1.45 by adding additional meglumine diatrizoate to the solution. The Axolotl used in this experiment are approved by MDIBL IACUC committee (no.# 20-04)

*Preparation of Ce3D-cleared mouse lymph node*

Lymph nodes were collected from a 13-week-old Male Ly5.1 mouse. Tissues were fixed for 24 hr at 4° C in 1 part fixative (Cat:554655, BD Biosciences) and 2 parts 1× PBS and incubated in blocking buffer for 4 days at 37°C. The buffer consisted of 30 mL Tris (Sigma-Aldrich #252859), 0.3 mL NMS (Sigma-Aldrich SML1128), 0.3 mL BSA (Sigma-Aldrich # A2058), and 0.09 mL TritonX100 (Sigma-Aldrich # T8787). Lymph nodes were stained for 3 days at 37°C while shaking in 500 μL blocking buffer, 2.5 μL SIRPa-cf594 [SIRPa Primary (BioLegend #144002)] conjugated using the CF™594 Mix-n-Stain™ Antibody Labeling Kit (BioLegend #92256), 2.5 μL CD11c-BV480 (Fisher Scientific #746392), 2.5 μL CD31-AF488 (BioLegend #102514) and 2.5 μL CD3-BV421 (BioLegend #100228). Tissues were then cleared with the Ce3D solution, consisting of 14 mL of 40% N-methyl-acetamide (Sigma-Aldrich # M26305), 25 μL Triton X-100 (Sigma-Aldrich # T8787), 20 g Histodenz (Sigma-Aldrich # D2158), and 125 μL Thioglycerol (Sigma-Aldrich #88640) for 1 day at room temperature. Procedures involving mice were approved by the Institutional Animal Care and Use Committee of the University of Washington in accordance with NIH guidelines.

*Preparation of Female p88 Slc17a7-IRES2-Cre;Ai14 cleared brain*

An adult (p100) Female p88 Slc17a7-IRES2-Cre;Ai14 mouse was transcardially perfused with 4% paraformaldehyde (PFA); the brain was extracted and post-fixed in 4% PFA for four hours prior to short-term storage in 1M PBS. The brain was then CUBIC-cleared using the same protocol as the MDA-231/OS-RC-2 brains with CUBIC-R+(N) to preserve GCaMP6s fluorescence.

*Preparation of SHIELD-cleared mouse embryo*

C57BL/6J (WT) and B6.129(Cg)-*Gt(ROSA)26Sor*$^{tm4(ACTB\text{-}tdTomato,\text{-}EGFP)Luo}$/J (“mTmG”) mice were maintained in a specific pathogen-free environment in the Animal Research and Care Facility (ARCF) at the University of Washington [31]. Mice were kept on a 12-h light/dark cycle with *ad libitum* access to food and water. Mice were originally bought from the Jackson Laboratory and bred in our facility. Female WT mice at least seven weeks of age were bred to male mTmG$^{+/+}$ mice; first day of plug positive was marked as embryonic day (E)0.5. On E12.5 or E13.5, dams were euthanized and $mTmG^{+/-}$ embryos were collected. Embryos underwent complete SHIELD tissue processing and clearing using equipment and reagents from Life Canvas Technologies following vendor-recommendations with the exception of halving sample volumes. Samples were kept in SHIELD perfusion solution for 1 day, SHIELD OFF solution for 1 day, SHIELD ON solution for 1 day, de-lipidated for 2-3 days, and incubated overnight in EasyClear 16-24 hours before imaging [32]. All procedures are approved by the University of Washington Institutional Animal Care and Use Committee.

*Preparation of ClearSee-processed Arabidopsis plant*

*AtFT::YFP-RCI2A* seedlings were used for imaging. *Arabidopsis* seedlings were grown on 1x Linsmaier and Skoog (LS) salt media (Caisson Laboratories) with 0.8% agarose. Plants were incubated for 14 days at 22°C under long day (16-hour light / 8-hour dark) conditions in which the red/far-red ratio was adjusted to 1 by supplementing far-red LED light. The seedlings were collected four hours after subjective dawn – for fixation [4% (w/v) paraformaldyhyde, in 1x PBS, pH 7.4] and subsequent clearing. The seedlings were fixed under -90kPA for 30 minutes and were then washed three times with 1x PBS. After final wash of 1x PBS, the solution was replaced with clearing solution. Clearing formulation is based on with minor modifications [9.3% (w/v) xylitol powder, 13.95% (w/v) sodium deoxycholate, and 23.25% (w/v) urea. This produces a solution with a refractive index of 1.41]. Samples were kept at room temperature in the dark, approximately two weeks for clearing. During the incubation, tubes were gently inverted every two days and the clearing solution replaced every week until the tissue had fully cleared.

*Preparation of EduClick-processed mouse prostate*

EdU (10mg/kg body weight) was administrated daily through i.p. injection to the adult mice for 3 days. After tissue dissection, the murine prostates were fixed in 10% neutrally buffered formalin at 4°C overnight, followed by washing in PBS at room temperature for three times. After permeabilization with 0.5% Triton X-100/PBS solution, the prostate tissues were incubated in the EdU staining solution from iClick™ EdU Andy Fluor 488 Imaging Kit (ABP Biosciences) for 3 days. Following the EdU staining and subsequent washing, a series of methanol/$H_2O$ solutions with a concentration gradient (20%, 40%, 60%, 80%, 100%) were used to dehydrate the prostates. Finally, samples were incubated in ethyl cinnamate at room temperature for clearing. All procedures are approved by the University of Washington Institutional Animal Care and Use Committee.

*Preparation of human prostate slice*

De-identified fresh prostate tissues were obtained prospectively from a genitourinary biorepository (approved by the University of Washington Institutional Review Board, IRB), with patient consent. The tissue was first placed in an individual 50 mL conical tube (Fisher Scientific #1443222) and stained in 70% ethanol at pH 4 with 1:200 dilution of Eosin-Y (Leica Biosystems #3801615) and 1:500 dilution of TO-PRO3 Iodide (Thermo-Fisher #T3605). The tissue was stained for 36 hr at room temperature with gentle agitation. Afterwards, the tissue was dehydrated in ethanol grades (25, 50, 75, 100%) for 8 hr per grade. The 100% grade was repeated to ensure removal of all water from the tissue. Finally, the tissue was cleared in ethyl-cinnamate (Sigma-Aldrich #112372) for 8 hr before imaging.

*Image processing of lymph nodes*

Image analysis of the cleared lymph node image was performed as described previously, with minor modifications [33]. Briefly, Imaris was used to segment cell objects, or to generate spots (i.e. central coordinates for each object). Next, a B220-Mask surface was created with the B220 channel using the parameters in Supplementary Table 3, and then the intensities were inverted. Next, spots were built from the B220 Mask and the CD31 channels, and surfaces built from the CD11c channel according to the respective creation parameters in Supplementary Table 3. Next, the position, volume, sphericity, and mean fluorescent intensity (MFI) of all channels, spots, and surface objects were exported from Imaris. The resulting .csv files were concatenated and imported into CytoMAP for analysis.

*CytoMAP analysis of lymph nodes*

CytoMAP version 1.4.20 was used for the analysis of spot and surface objects [33]. Once object statistics were imported into CytoMAP, the CD31 spots were clustered into 4 phenotypes using the Cluster Cells interface. The B220, SIRPa, CD31, CD11c, CD3, Sphericity, and Volume channels were standardized and used as clustering parameters. The CD31 clusters were next split into individual cell-type objects using the Annotate Clusters interface. The Region Statistics interface was used to plot the fold change of the MFI for each CD31 phenotype. Finally, phenotypes and cell densities were visualized using the New Figure interface. The positions of the spots were visualized both in 3D and as 100-μm stacks in 2D along the *z* axis.

| System | Hybrid OTLS | Blaze (Miltenyi Biotec) | SmartSPIM (Lifecanvas) | Alpha3 (PhaseView) | ct-dSPIM (ASI) | CTLS (3I) | MuVi SPIM CS (Luxendo) | Lightsheet 7 (Zeiss) |
|---|---|---|---|---|---|---|---|---|
| Illumination path | Single-sided | Double-sided | Double-sided | Double-sided | Single-sided | Double-sided | Double-sided | Double-sided |
| Collection path | Vertical (open-top) | Vertical | Vertical | Vertical | Vertical (inverted) | Vertical | Horizontal | Horizontal |
| Collection objective lenses | 2X (NA = 0.10)<br>24X (NA = 0.70) | 1.1X (NA = 0.10)<br>4X (NA = 0.35)<br>12X (NA = 0.53) | 3.6X (NA = 0.20)<br>15X (NA = 0.40) | 2X (NA = 0.14)<br>4X (NA = 0.28)<br>10X (NA = 0.50) | 16X (NA = 0.40)<br>24X (NA = 0.70) | 1X (NA = 0.20)<br>1.5X (NA = 0.37) | 10X (NA = 0.50)<br>20X (NA = 1.00) | 5X (NA = 0.16)<br>10X (NA = 0.50)<br>20X (NA = 1.00)<br>40X (NA = 1.00)<br>63X (NA = 1.00) |
| Horizontal FOV (mm) | 11.0 (2X)<br>1.0 (24X) | 33.0 (1.1X)<br>9.1 (4X)<br>3.0 (12X) | 3.7 (3.6X)<br>0.8 (15X) | 11.0 (2X)<br>5.5 (4X)<br>2.2 (10X) | 1.2 (16x)<br>1.0 (24x) | 22.0 (1X)<br>14.7 (1.5X) | 2.0 (10X)<br>1.0 (20X) | 4.0 (5X)<br>2.0 (10X)<br>1.0 (20X)<br>0.5 (40X)<br>0.3 (63X) |
| Axial resolution (μm) | 2.9 – 15.6 | 4.0 – 24.4 | 1.4 – 4.0 | 2.0 – 12.0 | >1.0 | 3.0 | 2.0 – 8.0 | 2.0 – 14.0 |
| Isotropic resolution? | No | No | No | No | No (single view)<br>Yes (dual view) | No | No (single view)<br>Yes (multi-view) | No |
| Refractive-index range | 1.33 – 1.56 | 1.33 – 1.56 | 1.33 – 1.56 | 1.33 – 1.56 | 1.33 – 1.56 | 1.33 – 1.56 | 1.33 – 1.51 (10X)<br>1.44 – 1.50 (20X) | 1.33 – 1.58 (5X)<br>1.33 (10X)<br>1.33, 1.38, 1.45, 1.53 (20X)<br>1.33 (40X)<br>1.33 (63X) |
| Maximum imaging depth (mm) | 10 | 17 (1.1X)<br>16 (4X)<br>10.9 (12X) | 12 | 15 | 5 (16X)<br>2 (24X) | 25 | 12 | 10 |
| Maximum specimen lateral size (X Y, mm) | Unconstrained | 24 x 50 | 20 x 25 (standard)<br>30 x 55 (extended) | 15 x 15 (standard)<br>15 x 25 (extended) | Unconstrained | 25 x 25 | 12 x 19 | 10 x 50 |

**Supplementary Table 1 | Commercially available cleared-tissue light-sheet microscopes.** The specifications of commercial light-sheet microscopes in comparison to our new hybrid OTLS system. For the collection path, "open-top" denotes that the objectives are below the specimen and enable unconstrained lateral imaging, whereas "inverted" denotes that the objectives are above the specimen, where unconstrained lateral imaging is also possible. Many of these specifications were obtained from the mesoSPIM publication [34].

| System | Hybrid OTLS | mesoSPIM V5 | ctASLM | LSTM | COLM |
|---|---|---|---|---|---|
| Illumination path | Single-sided | Double-sided | Single-sided | Double-sided | Dual-sided |
| Collection path | Vertical (open-top) | Horizontal | Horizontal | Vertical (inverted) | Vertical |
| Collection objective lenses | 2X (NA = 0.10)<br>24X (NA = 0.70) | 1X (NA = 0.25) | 16X (NA = 0.40)<br>24X (NA = 0.70) | 10X (NA = 0.60)<br>25X (NA = 1.00) | 10x (NA = 0.60)<br>25x (NA = 1.00) |
| Horizontal FOV (mm) | 11.0 (2X)<br>1.0 (24X) | 20.0 | 1.0 (16X)<br>1.2 (24X) | 2.0 (10X)<br>0.8 (25X) | 2.0 (10X)<br>0.8 (25X) |
| Axial resolution (μm) | 2.9 – 15.6 | 6.5 | 0.3 – 0.6 | 4.0 – 11.0 | 1.2 |
| Isotropic resolution? | No | Yes (ASLM) | Yes (ASLM) | No | No |
| Refractive-index range | 1.33 – 1.56 | 1.33 – 1.56 | 1.33 – 1.56 | 1.33 – 1.52 (10X)<br>1.41 – 1.52 (25X) | 1.33 – 1.52 (10X)<br>1.41 – 1.52 (25X) |
| Maximum imaging depth (mm) | 10 | 52 | 12 (16x)<br>10 (24x) | 8 | 8 |
| Maximum specimen lateral size (X Y, mm) | Unconstrained | 52 x 100 | 12 x 12 (16x)<br>10 x 10 (24x) | Unconstrained | 45 x 45 |

**Supplementary Table 2 | Selected list of academic cleared-tissue light-sheet microscopes.** The specifications of a selected list of academic light-sheet microscopes in comparison to our new hybrid OTLS system [24, 34-36]. For the collection path, "open-top" denotes that the objectives are below the specimen and enable unconstrained lateral imaging, whereas "inverted" denotes that the objectives are above the specimen, where unconstrained lateral imaging is also possible. ASLM: axially swept light-sheet microscopy.

| | Radius | Surface detail | Background subtraction | Channel threshold | Quality threshold | Splitting seed diameter | Volume threshold |
|---|---|---|---|---|---|---|---|
| **B5220 mask** | N/A | 3 μm | None | 2.56 | None | None | None |
| **B220 spots** | 5 | N/A | True | N/A | <1.5<br>>25 | N/A | N/A |
| **CD31 spots** | 5 | N/A | True | N/A | 4 | N/A | N/A |
| **CD11c surfaces** | N/A | 1 μm | 15 μm | 0.5 | 0.8 | 10 μm | None |

**Supplementary Table 3 | CytoMAP parameters.** Surface-creation parameters used during image processing of the lymph-node dataset.

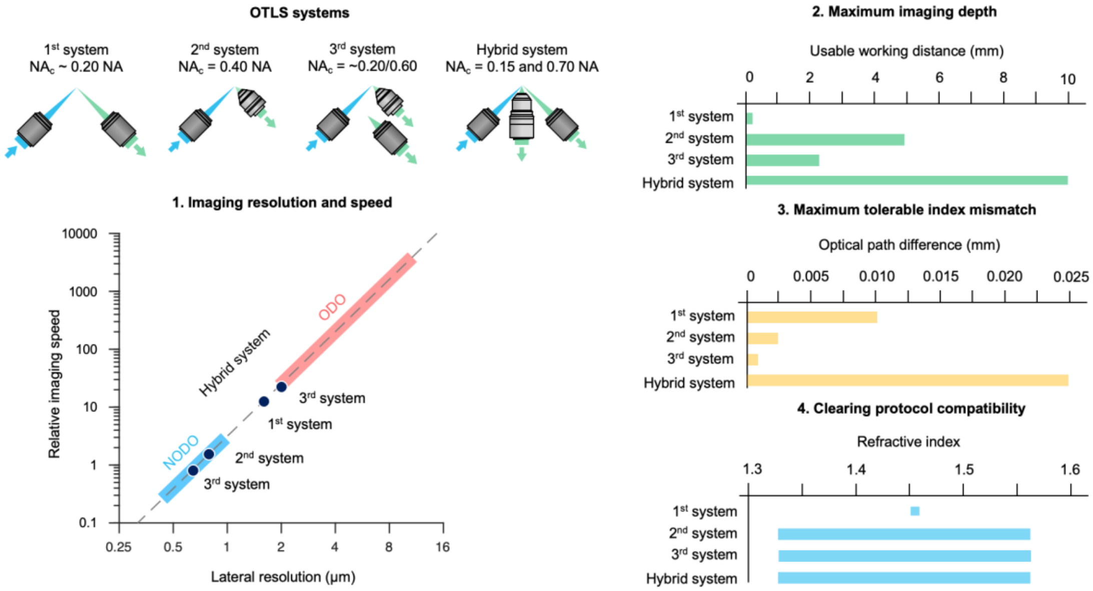


**Supplementary Figure 1 | Comparison of new and past OTLS microscope configurations.** The specifications (imaging resolution, speed, imaging depth, index-mismatch sensitivity, and clearing-protocol compatibility) of our new hybrid OTLS system as compared to three previous generations of OTLS microscopes [1, 2, 4]. Note, in the imaging resolution and speed panel, the NODO and ODO paths are shown over the range corresponding to either NA-maximized or FOV-maximized imaging. Our previous systems could also be reconfigured over a certain range, but are plotted here as single data points corresponding to the published version of each system.

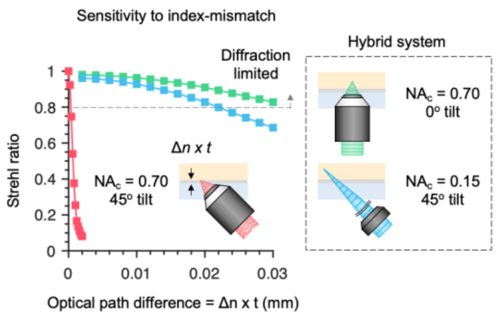


**Supplementary Figure 2 | Comparison between NODO and single-objective light-sheet architectures.** Unlike ODO, which is extremely sensitive to refractive-index mismatch at higher NAs (red curve), NODO uses a collection objective oriented in the normal (vertical) direction, which is much less sensitive to index mismatch (green curve). The index-mismatch sensitivity for the low-resolution ODO collection path of the hybrid OTLS system is also shown (blue curve). Note that the index mismatch is represented by the optical path difference, Δ*n* x *t*, where Δ*n* is the refractive-index difference between the holder material and the immersion medium/tissue, and *t* is the thickness of the sample holder. A Strehl ratio of >0.8 is often considered as ideal “diffraction-limited” imaging.

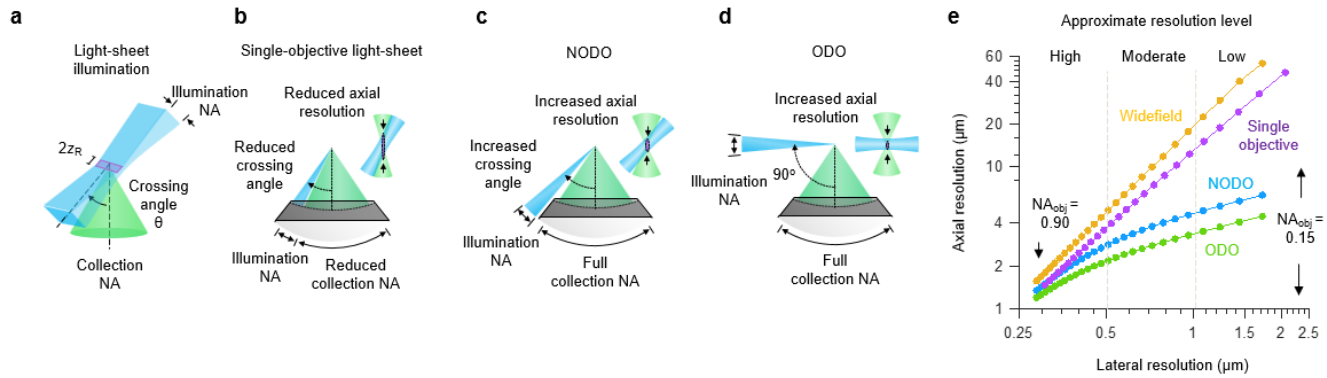


| Illumination NA | Single-objective light-sheet | NODO | ODO | Simulation inputs |
|---|---|---|---|---|
| FOV = 2048 x 0.61 x λ / ($NA_{obj}$) / 2 | θ = $\sin^{-1}(NA_{obj} / n) - \sin^{-1}(NA_i / n)$ | θ = 45 deg. | θ = 90 deg. | λ = 0.5 μm |
| $2z_R$ = FOV / 8 | $NA_c = NA_{obj} – NA_i / 2$ | $NA_c = NA_{obj}$ | $NA_c = NA_{obj}$ | $n$ = 1.5 |
| $NA_i = z_R$ x λ / π / $n$ | | | | $NA_{obj}$ = 0.15 – 0.9 (0.03 steps) |

**Supplementary Figure 3 | Extended comparison of single-objective, NODO, and ODO optical performance.** **(a)** For a given objective NA, the FOV at Nyquist sampling was calculated for a 2048 x 2048 pixel sCMOS camera. The illumination NA was then chosen such that the Rayleigh length, $2z_R$, would fill 1/8 of the chip (256 pixels). This is a reasonable vertical height for systems that perform stage-scanned imaging, such as the current hybrid OTLS system. **(b)** For the single-objective light-sheet architecture, there is a coupled relationship between the illumination NA, collection NA, and crossing angle of the two beams. **(c)** For the NODO architecture, the paths are decoupled. **(d)** For the ODO architecture, the paths are decoupled and orthogonal to one another. **(e)** Simulation results for the lateral and axial resolution of the single-objective light-sheet and NODO architectures for objective NA's ($NA_{obj}$) ranging from 0.15 – 0.90. At low and moderate objective NAs, there is a substantial performance benefit of NODO vs. single-objective light-sheet, approaching that of ODO (but with superior imaging depth and tolerance to index-mismatch). The optical performance for widefield microscopy is also shown.

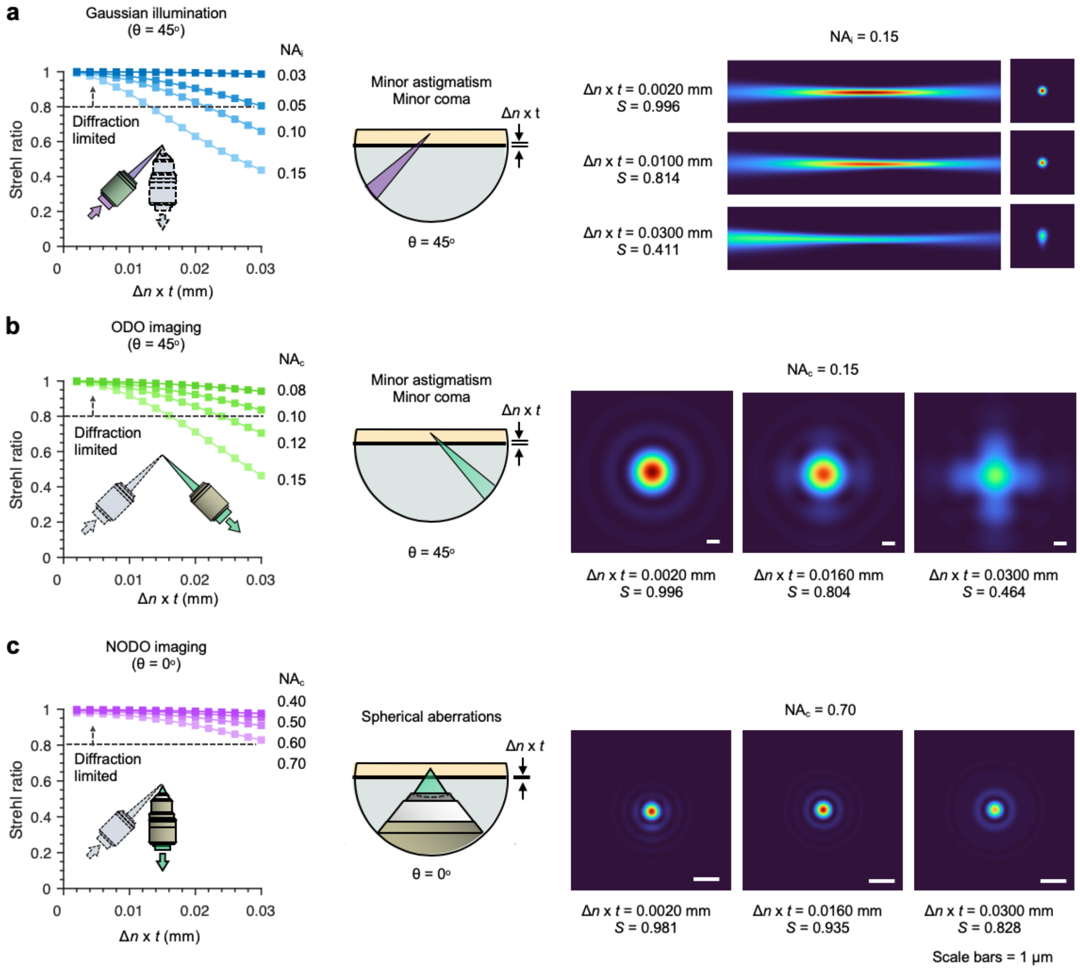


**Supplementary Figure 4 | Index-mismatch sensitivity of the hybrid OTLS system.** ZEMAX simulations of the index-mismatch sensitivity for all three optical paths in the hybrid OTLS system. The index mismatch is given by the optical path difference, Δ*n* x *t*, where Δ*n* is the refractive-index difference between the holder and immersion medium/tissue, and *t* is the thickness of the holder. **(a)** Results for a Gaussian illumination light sheet with corresponding cross-sectional irradiance profiles. The angle of illumination is 45 deg **(b)** Results for the ODO collection path, with corresponding point spread functions. The angle of collection is also 45 deg **(c)** Results for the NODO collection path, with corresponding point spread functions. The angle of collection is 0 deg. In **(a-c)**, $NA_i$ refers to illumination NA, and $NA_c$ refers to collection NA. Note that a Strehl ratio of >0.8 is often considered as ideal (“diffraction-limited” imaging).

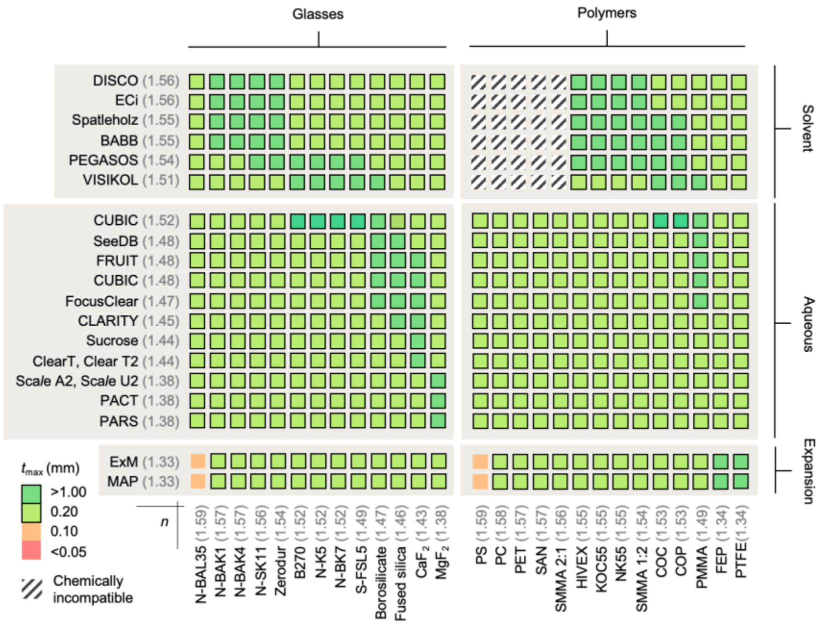


**Supplementary Figure 5 | Compatible combinations of clearing protocols and sample-holder materials.** Based on the ZEMAX index-mismatch simulations, a matrix to illustrate clearing protocols vs. compatible holder materials can be generated. Based on the maximum index mismatch, the corresponding maximum specimen holder thickness, $t_{max}$ is calculated. Thickness values greater than 0.20 mm are considered to be practically feasible. The hybrid OTLS system is largely insensitive to index mismatch, enabling the combination of virtually any clearing protocol and specimen-holder material. Note that the latest CUBIC clearing protocol uses a refractive index of 1.52, which is also compatible with the glasses and plastics shown in the figure.

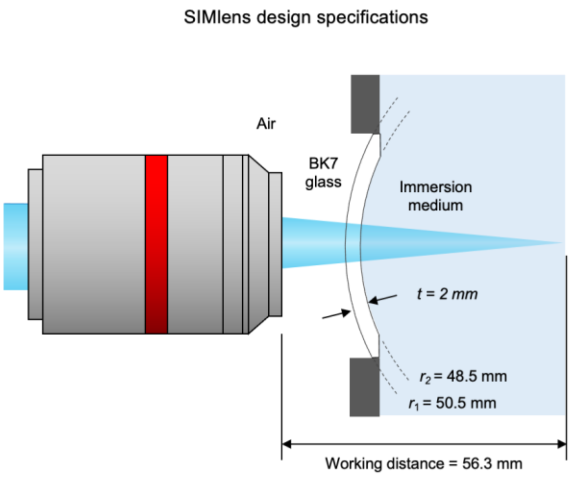


**Supplementary Figure 6 | SIMlens specifications.** The specifications of the SIMlens used on the illumination and ODO optical paths are shown. The lens is fabricated of BK7 glass and is 2-mm thick. The radii of curvature of the lens surfaces are 48.5 and 50.5 mm respectively (identical center of curvature). The lens is positioned such that the center of curvature for both lens surfaces is coincident with the on-axis focus of the objective. This causes all rays traveling to or from this focal point to transition between the SIMlens interfaces at a normal (90-deg) angle where refraction is negligible, thereby suppressing any aberrations.

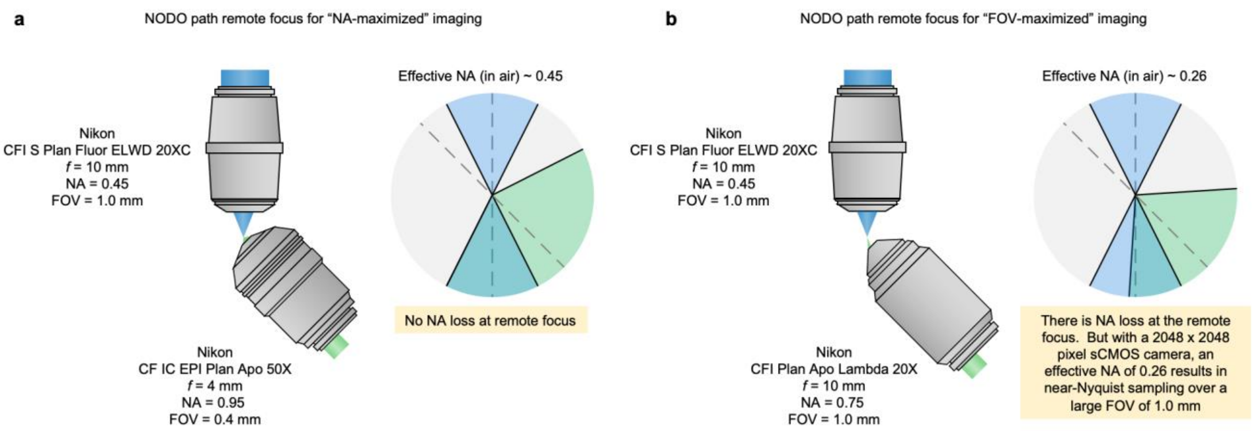


**Supplementary Figure 7 | Remote-focus layout for NODO imaging.** Our NODO imaging architecture enables the use of commercial air objectives at the remote focus, as opposed to bespoke objective assemblies. **(a)** Using a crossing angle of 45 deg, the effective NA of the NODO imaging path is a lossless ~0.45 (in air) when using a 50x 0.95 NA air objective. This configuration provides NA-maximized imaging **(b)** By switching to a 20x 0.75 NA remote objective, the effective NA is reduced to ~0.26 (in air). Despite the loss in effective NA, Nyquist sampling at this NA on a standard 2048 x 2048 pixel sCMOS camera yields a field of view of ~ 1.0 mm. Therefore, this configuration provides FOV-maximized imaging.

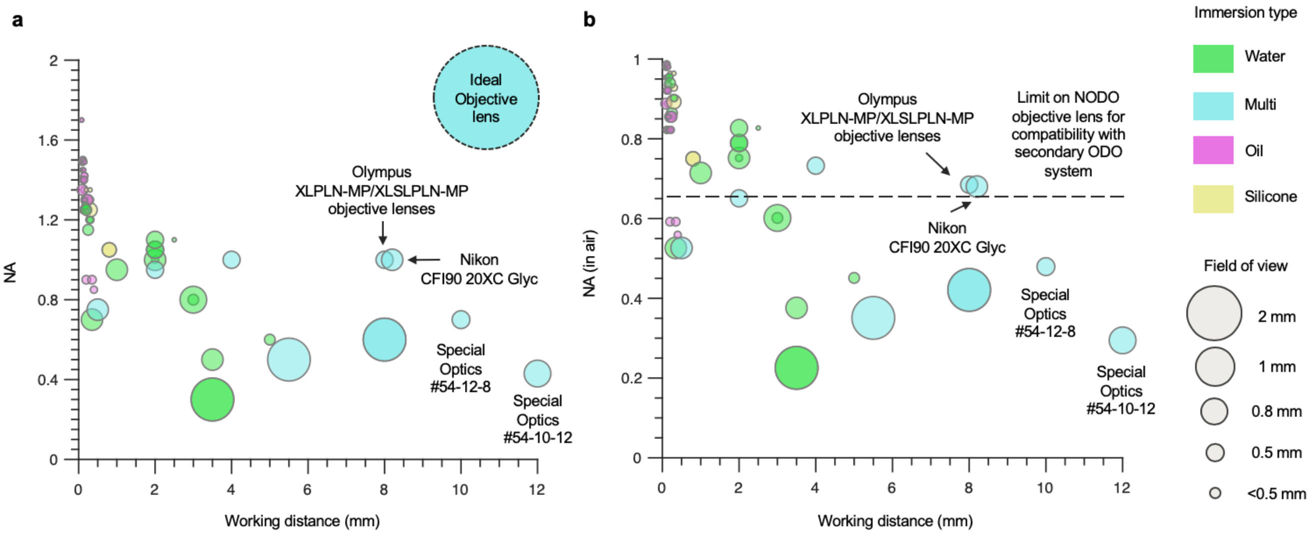


**Supplementary Figure 8 | Candidate primary microscope objectives (O1) for NODO imaging. (a)** Plot summarizing objectives currently available from Olympus, Nikon, and Special Optics for NODO imaging of cleared tissues. The color of each data point denotes the immersion type (water, multi, oil, or silicone) and the size of the point denotes the field of view. As can be seen, in current objectives, there is a distinct trade-off between magnification, NA, and working distance. An ideal objective, with a large field of view, high NA, and long working distance is also plotted. The multi-immersion objective (Special Optics #54-12-8) used in the current system's NODO imaging path is also highlighted. **(b)** The same data is plotted but with the y axis showing the NA in air as opposed to the immersion medium. In this plot, a line is drawn above which hybrid imaging would no longer be possible, as the optical cone angle of the NODO imaging objective would surpass 45 deg (NA > 0.65 in air), thereby preventing additional optical paths at 45 deg for ODO imaging.  For example, the Olympus XLPLN/XLSPLN and Nikon CFI90 20XC Glyc objectives would be attractive for NODO imaging but would not allow for multi-scale imaging with a separate ODO imaging path due to its large NA. Note, the level of chromatic aberration correction for each lens is not indicated in either plot. A data file containing the information used to generate the plots is available as **Supplementary Data**.

**a**

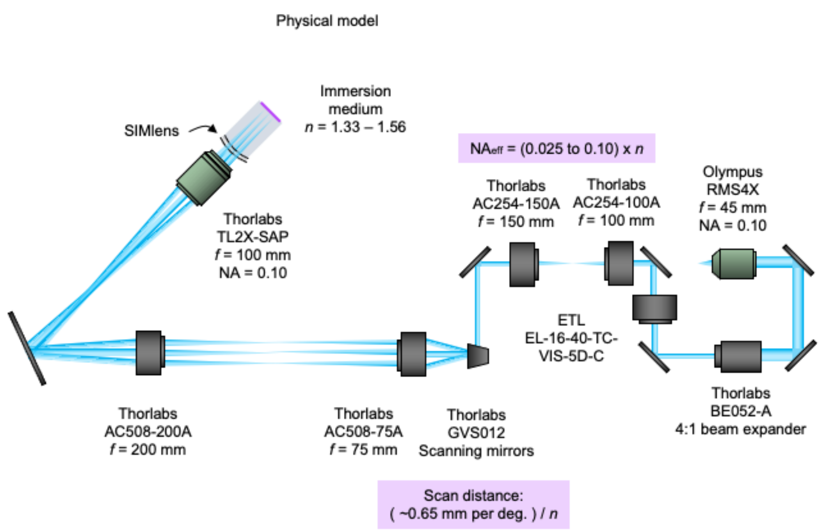


**c**

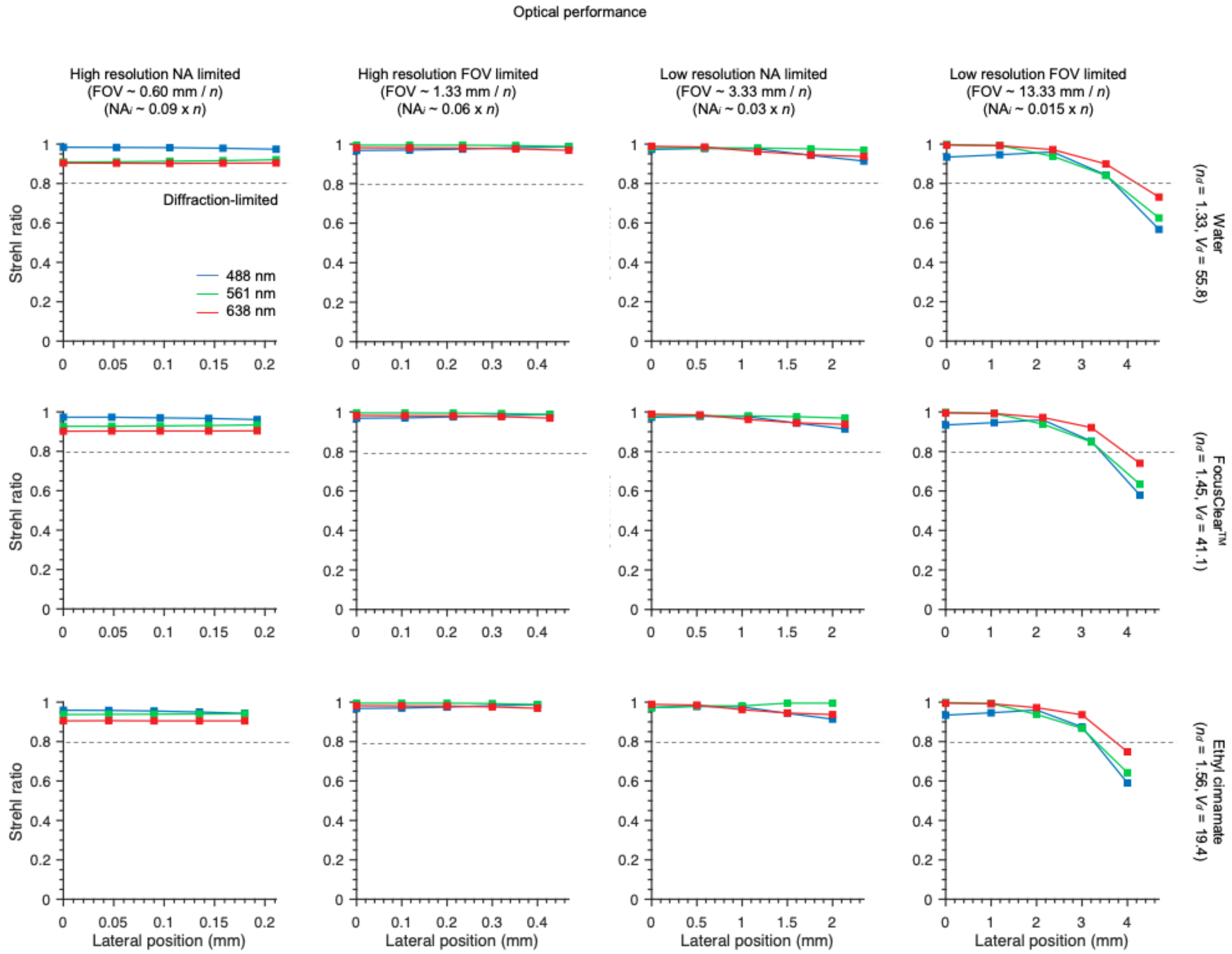


**Supplementary Figure 9 | Illumination optics models. (a)** The physical model of the hybrid OTLS system illumination optics is shown. Light is collimated using a 20X objective and passed through a 4:1 variable beam expander. The adjusted beam is then scanned using a pair of galvanometric scanning mirrors, which is then relayed to the back focal plane of the 2X illumination objective using a pair of relay lenses. Finally, the light passes through the SIMlens and into the specimen. The effective illumination NA can be varied from ~ 0.025 – 0.10, multiplied by the refractive-index of the immersion medium, *n*. The width of the light sheet can similarly be tuned using one of the scanning mirrors, which provides a scanning distance of ~0.65 mm / deg, divided *n*. The second scanning mirror is used to align the light sheet to the focal planes of the collection objectives. **(b)** Results for the Strehl ratio as a function of lateral position across the width of the sheet. The four columns correspond to the four imaging modes, high-resolution (NODO) NA-maximized and FOV- maximized, and low-resolution (ODO) NA-maximized and FOV- maximized. The three rows correspond to different immersion media, water (*n* ~ 1.33), FocusClear™ (*n* ~ 1.46), and ethyl cinnamate (*n* ~ 1.56). The colored lines on each plot denote wavelength, 488 nm (blue), 561 nm (green), and 638 nm (red).

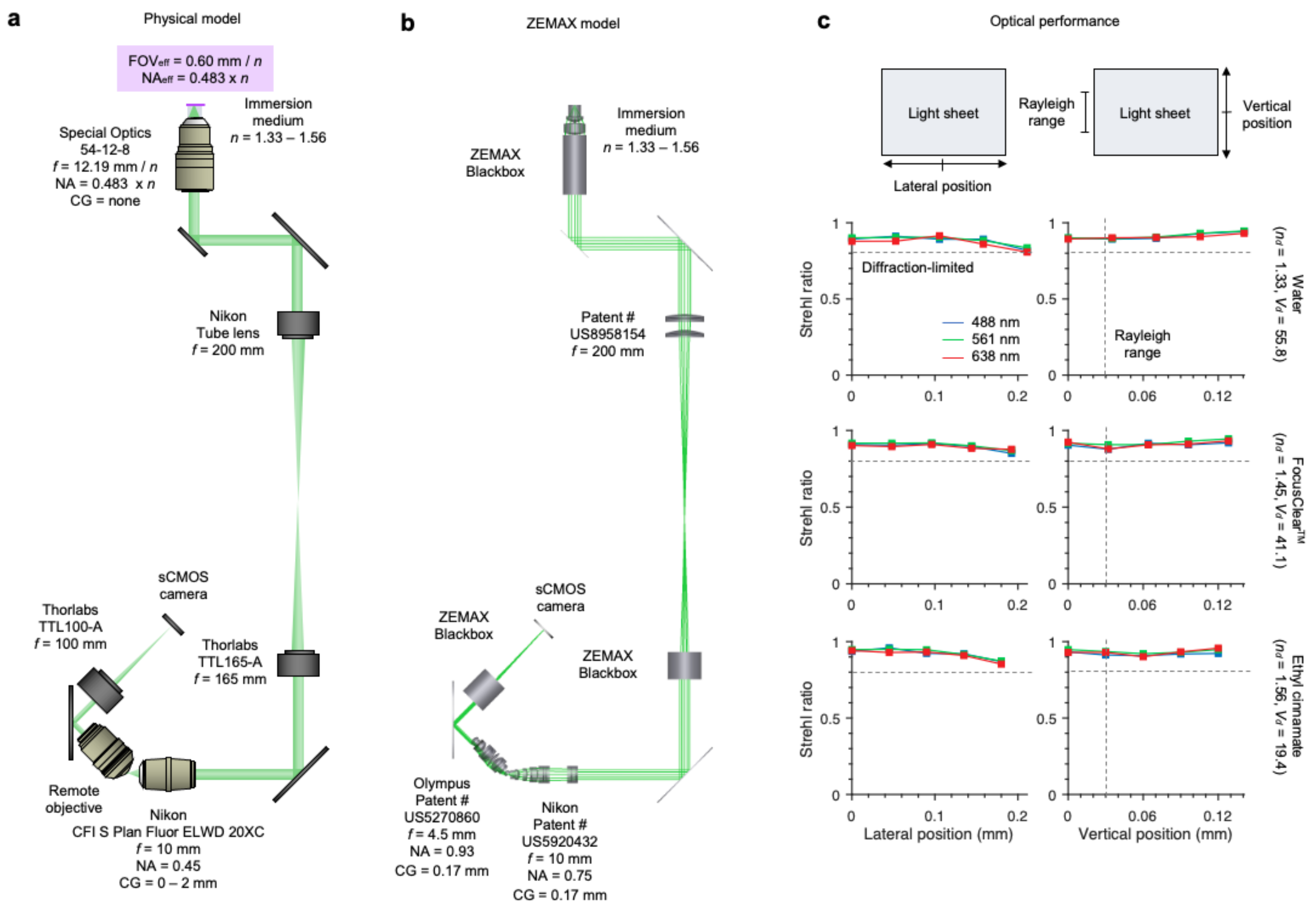


**d**

Remote objective options for NA-maximized NODO imaging

| Company | Model # | Cover glass | NA | FOV | $NA_{eff}$ | $M_{total}$ | Nyquist rate |
|---|---|---|---|---|---|---|---|
| Nikon | CFI Plan Apo Lambda 40XC | 0.11 – 0.23 mm | 0.95 | 0.5 mm | 0.45 x *n* | 20.00 x *n* | 2.17 |
| Nikon | CF IC EPI Plan Apo 50X | None | 0.95 | 0.4 mm | 0.45 x *n* | 25.00 x *n* | 2.71 |
| Olympus | UPLSAPO40X2 | 0.11 – 0.23 mm | 0.95 | 0.5 mm | 0.45 x *n* | 22.22 x *n* | 2.41 |
| Olympus | MPLAPON50X | None | 0.95 | 0.4 mm | 0.45 x *n* | 27.78 x *n* | 3.01 |

**Supplementary Figure 10 | NA-maximized NODO collection optics model**. **(a)** The physical model of the NA-limited NODO collection optics within the hybrid OTLS system. Fluorescence is collected by a multi-immersion objective (*f* = 12.19 / *n*) and relayed to a remote focus using tube lenses (*f* = 200 and 165 mm), and a 20X objective (*f* = 10 mm). For aberration free imaging, the total magnification from the specimen to the remote focus should be equal to *n*. For the optics chosen, the total magnification is extremely close: ~0.994 *n*. The remote imaging arm is tilted by 45 deg and uses a remote objective and tube lens (*f* = 100 mm) to image the remote focus onto a sCMOS camera. The effective numerical aperture of the multi-immersion objective is 0.45 x *n*, with an effective FOV of 0.60 mm / *n*. **(b)** Corresponding ZEMAX model. Lens prescriptions or blackbox files were available for all components, except for the remote imaging objectives. For these objectives, US Patent # 5920432 and #5270860 were used. **(c)** Results for the Strehl ratio as a function of lateral position across the width of the sheet (left) and along the length of the sheet (right). The three rows correspond to different immersion media, water (*n* ~ 1.33), FocusClear$^{TM}$ (*n* ~ 1.46), and ethyl cinnamate (*n* ~ 1.56). The colored lines on each plot denote wavelength, 488 nm (blue), 561 nm (green), and 638 nm (red). **(d)** Various options for the remote objective, and the corresponding imaging specifications.

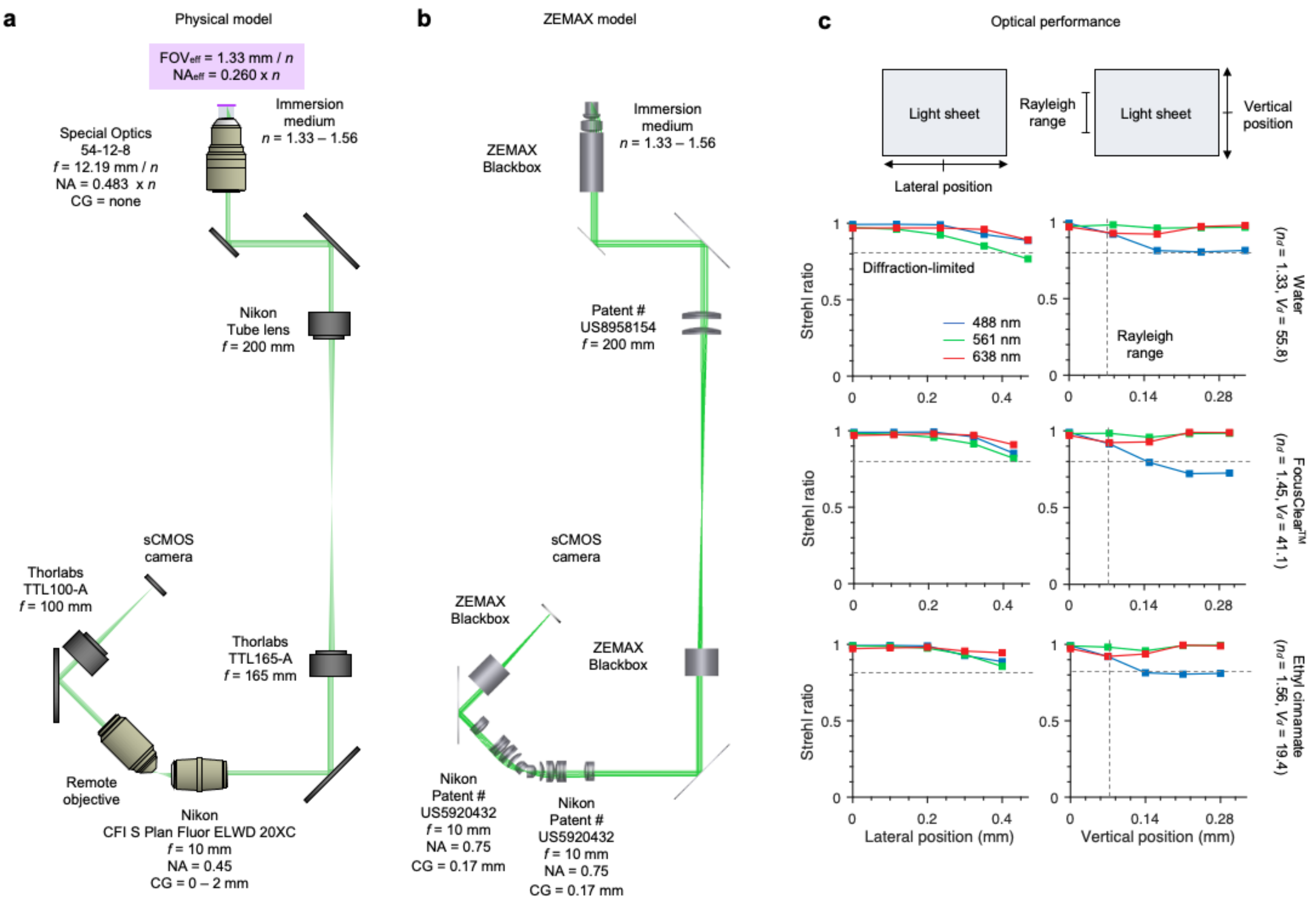


**d**

Remote objective options for FOV-maximized NODO imaging

| Company | Model # | Cover glass | NA | FOV | $NA_{eff}$ | $M_{total}$ | Nyquist rate |
|---|---|---|---|---|---|---|---|
| Nikon | CFI Plan Apo Lambda 20X | 0.17 mm | 0.75 | 1 mm | ~0.26 x *n* | 10.00 x *n* | 2.02 |
| Nikon | CFI Super Fluor 20X | 0.17 mm | 0.75 | 1 mm | ~0.26 x *n* | 10.00 x *n* | 2.02 |
| Nikon | CFI S Plan Fluor LWD 20XC | 0 – 1.8 mm | 0.70 | 1 mm | ~0.22 x *n* | 10.00 x *n* | 2.39 |
| Olympus | UPLSAPO20X | 0.17 mm | 0.75 | 1 mm | ~0.26 x *n* | 11.11 x *n* | 2.24 |
| Olympus | UPLXAPO20X | 0.17 mm | 0.80 | 1 mm | ~0.29 x *n* | 11.11 x *n* | 2.01 |
| Olympus | UCPLFLN20X | 0 – 1.6 mm | 0.70 | 1 mm | ~0.22 x *n* | 11.11 x *n* | 2.65 |

**Supplementary Figure 11 | FOV-maximized NODO collection optics model. (a)** The physical model of the FOV-limited NODO collection optics in the hybrid OTLS system. The optics are identical to **Supplementary Figure 10**, except the imaging arm now uses a 20X remote air objective and the same tube lens ($f$ = 100 mm) to image the remote focus onto a sCMOS camera. **(b)** Corresponding ZEMAX model. Lens prescriptions or blackbox files were available for all components, except for the remote imaging objectives. For both objectives, US Patent # 5920432 was used. **(c)** Results for the Strehl ratio as a function of lateral position across the width of the sheet (left) and along the length of the sheet (right). **(d)** Various options for the remote objective, and the corresponding imaging specifications.

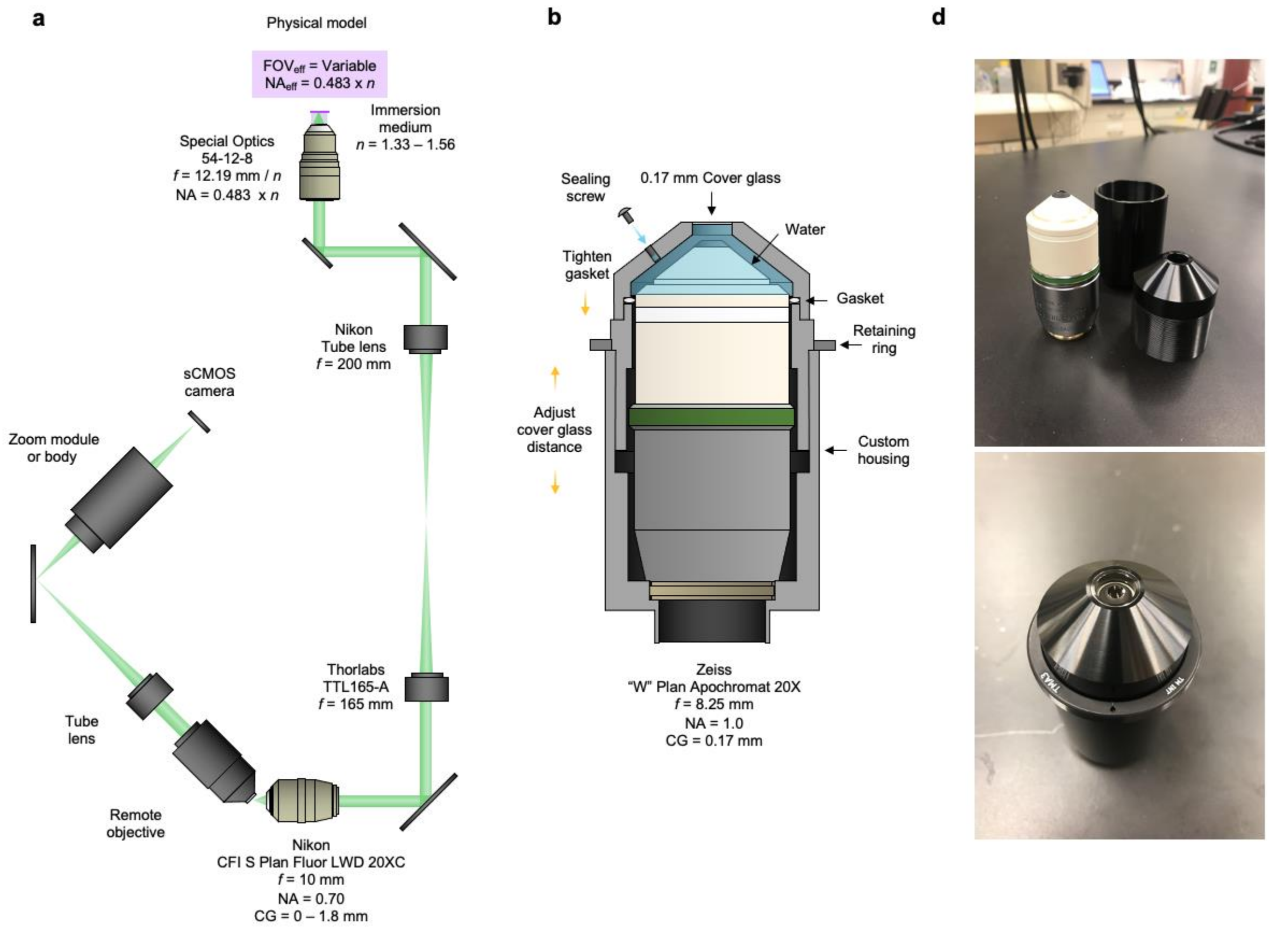


**e** Remote objective options for NA-maximized NODO imaging

| Company | Model # | Cover glass | NA | FOV | NA$_{eff}$ | M$_{total}$ | Nyquist rate |
|---|---|---|---|---|---|---|---|
| Zeiss | "W" Plan Apochromat 20X | 0.17 mm | 1.0 | 1 mm | 0.483 x $n$ | Variable | Variable |
| Olympus | XLUMPLFLN20XW | None | 1.0 | 1 mm | 0.483 x $n$ | Variable | Variable |
| Special Optics | AMS-AGY v1.0 | None | 1.0 | 0.15 – 0.25 mm | 0.483 x $n$ | Variable | Variable |
| Special Optics | AMS-AGY v2.0 | None | 1.0 | 0.45 – 0.75 mm | 0.483 x $n$ | Variable | Variable |

**Supplementary Figure 12 | Alternative NODO collection optics model. (a)** The physical model of alternative NODO collection optics is shown. Fluorescence is collected by a multi-immersion objective ($f$ = 12.19 / $n$) and relayed to a remote focus using tube lenses ($f$ = 200 and 165 mm), and a 20X objective ($f$ = 10 mm). The remote imaging arm is tilted by 45 deg and uses a remote objective and tube lens ($f$ = 100 mm) to image the remote focus onto a sCMOS camera. By using a liquid or solid immersion remote objective, the full NA is maintained, and the effective NA of the system is 0.483 x $n$ **(b)** Custom designed sleeve to convert a Zeiss "W" Plan Apochromat 20X objective into a remote objective assembly. **(c)** Results for the fabricated sleeve shown in **(b)**. **(d)** Various options for the remote objective, and the corresponding imaging specifications.

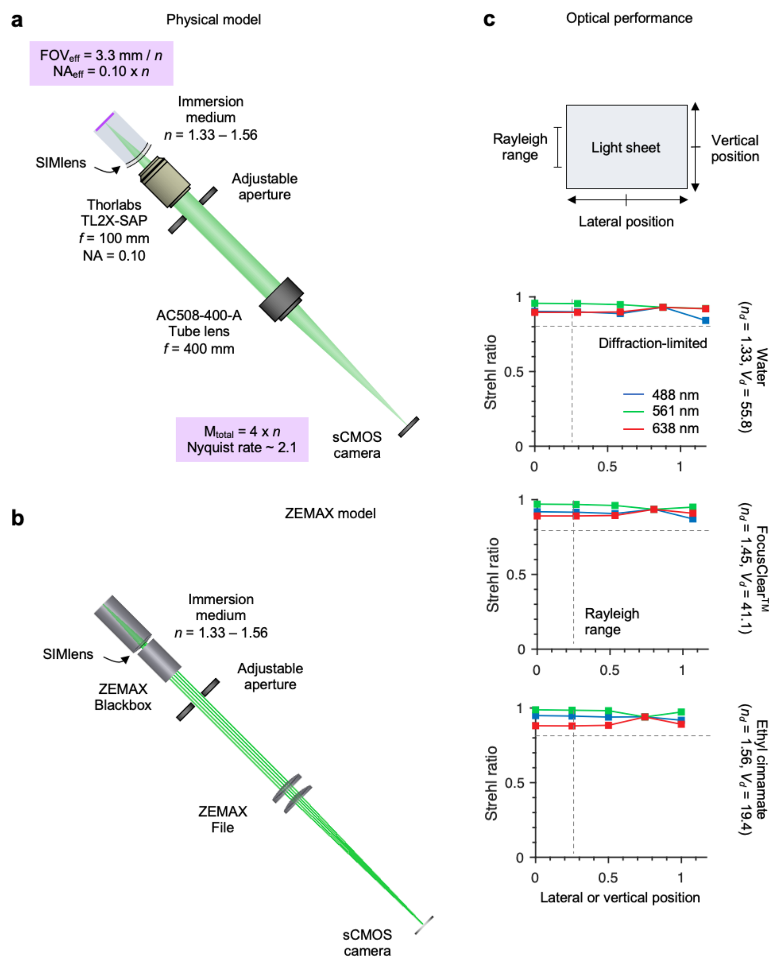


**Supplementary Figure 13 | NA-maximized ODO collection optics model. (a)** The physical model of the NA-limited ODO collection optics within a hybrid OTLS system. Fluorescence is collected by a 2X (NA = 0.10) air objective. The objective is combined with the SIMlens, which provides multi-immersion compatibility, and increases the numerical aperture of the objective by *n*. Light is focused onto the sCMOS camera using a long focal length tube lens (*f* = 400 mm). The effective numerical aperture of the SIMlens and air objective is 0.10 x *n*, with an effective FOV of 3.33 mm / *n*. The total magnification of the imaging arm is 4 x *n*, which yields a Nyquist sampling rate of ~ 2.1 on the sCMOS camera. **(b)** Corresponding ZEMAX model. Lens prescriptions or blackbox files were available for all components. **(c)** Results for the Strehl ratio as a function of lateral position across the width of the sheet or along the length of the sheet. Unlike the NODO collection optics, where the two directions are modeled independently due to the tilted remote focus, the two directions are identical and symmetric for the ODO imaging path.

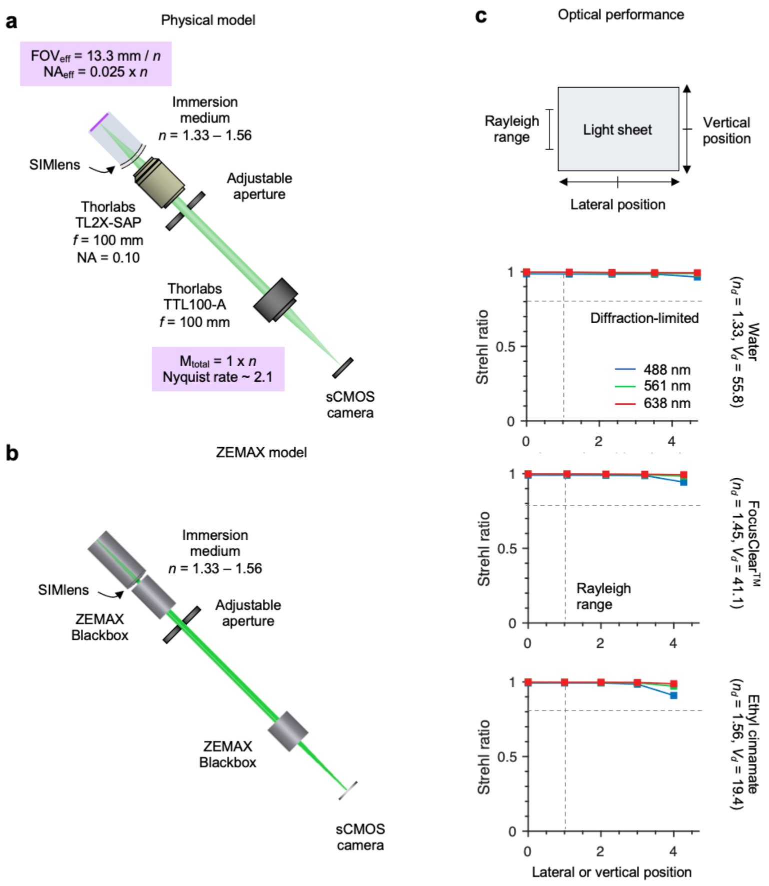


**Supplementary Figure 14 | FOV-maximized ODO collection optics model. (a)** The physical model of the FOV-limited ODO collection optics in the hybrid OTLS system. Fluorescence is collected by a 2X (NA = 0.10) air objective. The objective is combined with a SIMlens, which provides multi-immersion compatibility, and increases the numerical aperture of the objective by *n*. Light is focused onto the sCMOS camera using a short focal length tube lens (*f* = 100 mm). The effective numerical aperture of the SIMlens and air objective is 0.025 x *n*, with an effective FOV of 13.33 mm / *n*. The total magnification of the imaging arm is 1 x *n*, which yields a Nyquist sampling rate of ~ 2.1 on the sCMOS camera. **(b)** Corresponding ZEMAX model. Lens prescriptions or blackbox files were available for all components. **(c)** Results for the Strehl ratio as a function of lateral position across the width of the sheet or along the length of the sheet. Unlike the NODO collection optics, where the two directions are modeled independently due to the tilted remote focus, the two directions are identical and symmetric for the ODO imaging path.

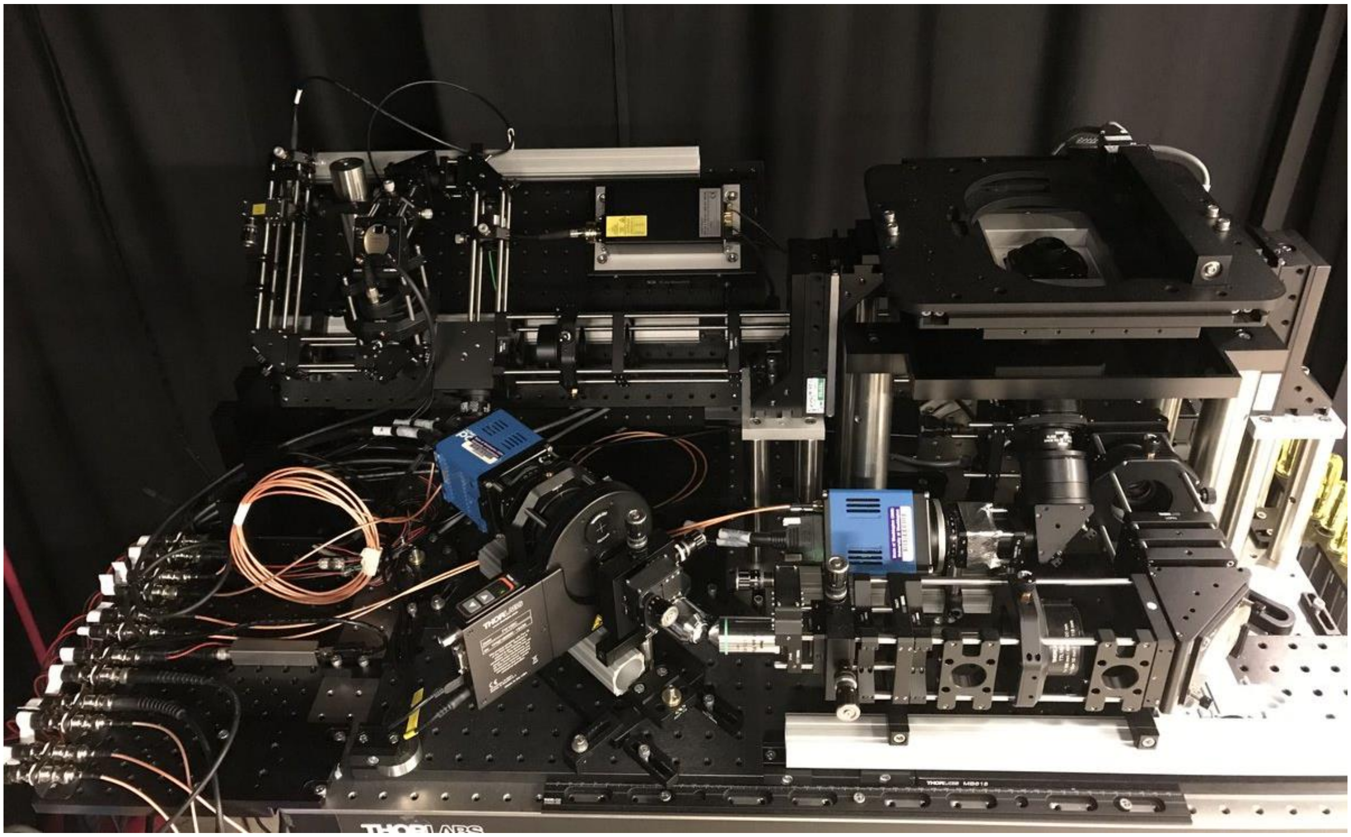

**Supplementary Figure 15 | Hybrid OTLS system in practice.** Photograph of the hybrid OTLS system. The illumination optics are housed on a single 18 x 36 in. breadboard angled at 45 deg. The NODO and ODO optics are placed on flat breadboards. The NODO imaging optics are housed on a series of connected breadboards, which can be translated in one direction to adjust the distance between the imaging optics and the back focal plane (BFP) of the multi-immersion objective. This enables fine alignment when switching the immersion medium, as the axial position of the BFP of the objective is a function of refractive-index, $n$. The position of the BFP relative to the flange of the objective is approximately -11.75 mm - 22.90 mm x $n$ + 5.87 mm x $n^2$. Finally, the remote objective, final tube lens, and sCMOS camera are all positioned atop another breadboard, which can be translated and rotated to align the remote imaging arm with respect to the remote focus.

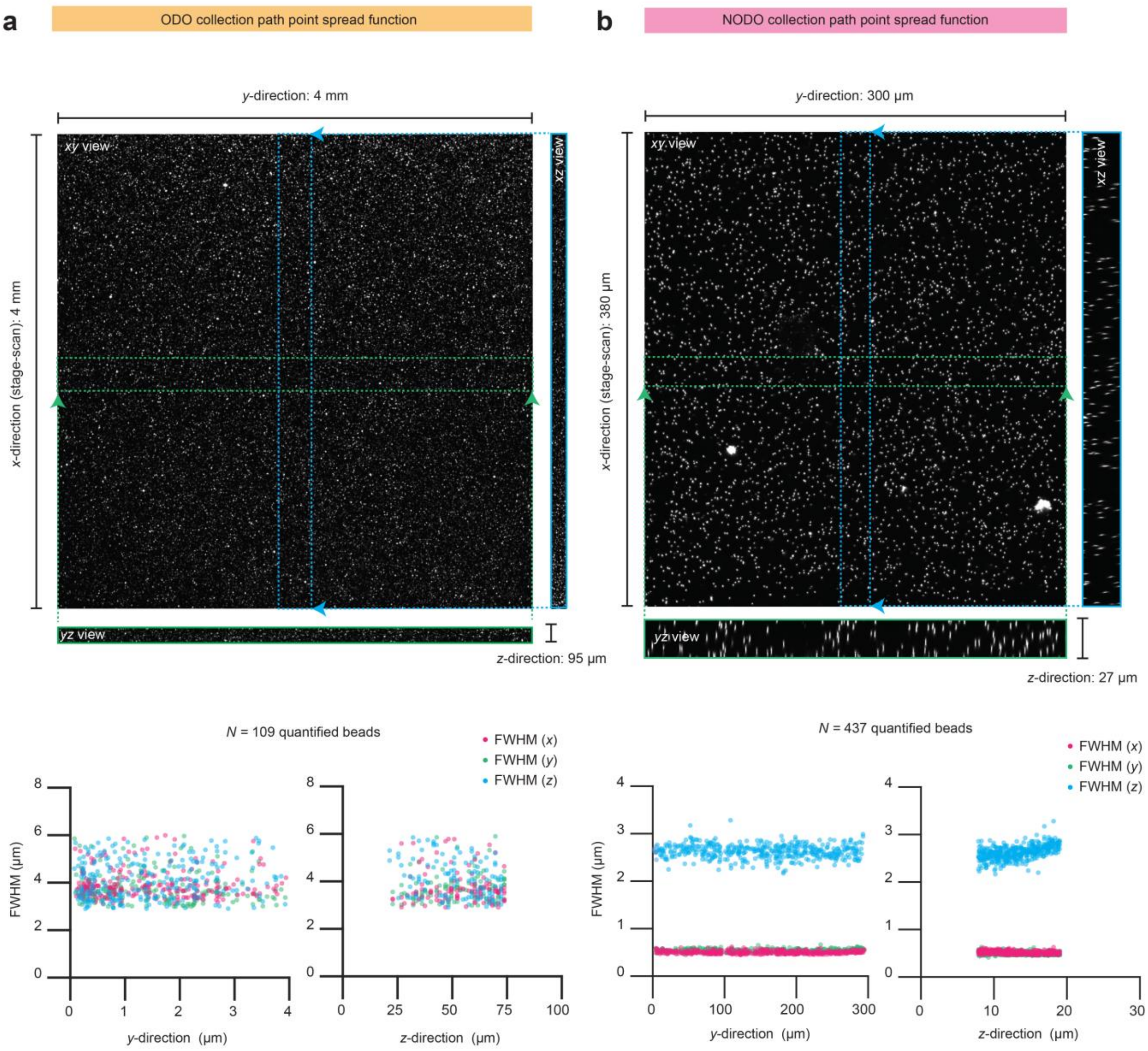


**Supplementary Figure 16 | Point spread function measurements.** Point spread function measurements for the ODO **(a)** and NODO **(b)** imaging paths. Measurements were taken from gold nanoparticles (150-nm diameter) embedded in 1% v/v agarose. The agarose phantom was cleared using ECi ($n$ = 1.56) and reflectance from the beads measured using 638 nm illumination. For both datasets, the XYZ centers of the beads were identified. For the ODO dataset, only beads with a clear 16 x 16 x 16 µm volume were considered, resulting in a total of $N$ = 109 beads. For the NODO dataset, only beads with a clear 2 x 2 x 12 µm were considered, resulting in a total of $N$ = 437 beads. Each bead was fit to a 3D Gaussian distribution to quantify the full-width half-maximum (FWHM) value of each bead in the XYZ directions. The resulting values are plotted in graphs versus the y-direction (lateral across the camera chip) and z-direction (vertical across the camera chip). Results are not plotted as a function of the x-direction (stage scanning) as there is no anticipated difference in resolution as a function of stage-scanning position ($x$) within the bead phantom.

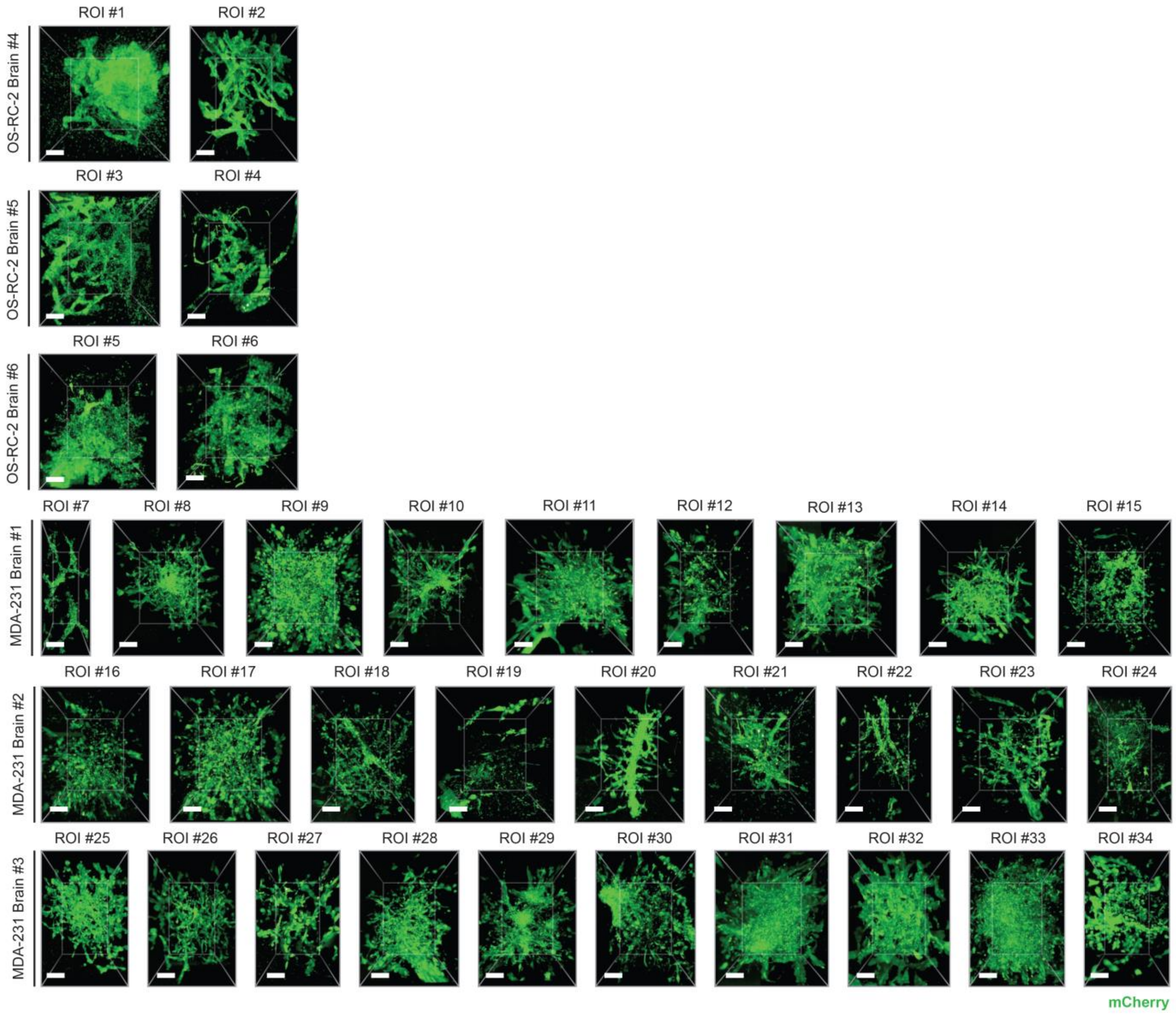


**Supplementary Figure 17 |** Regions of interests from metastatic brain specimens. All *N* = 34 metastatic colonies collected from the metastatic brain specimens (3 brains from OS-RC-2 cancer cell line, 3 brains from MDA-231 cancer cell line). All scale bars represent 10 μm.

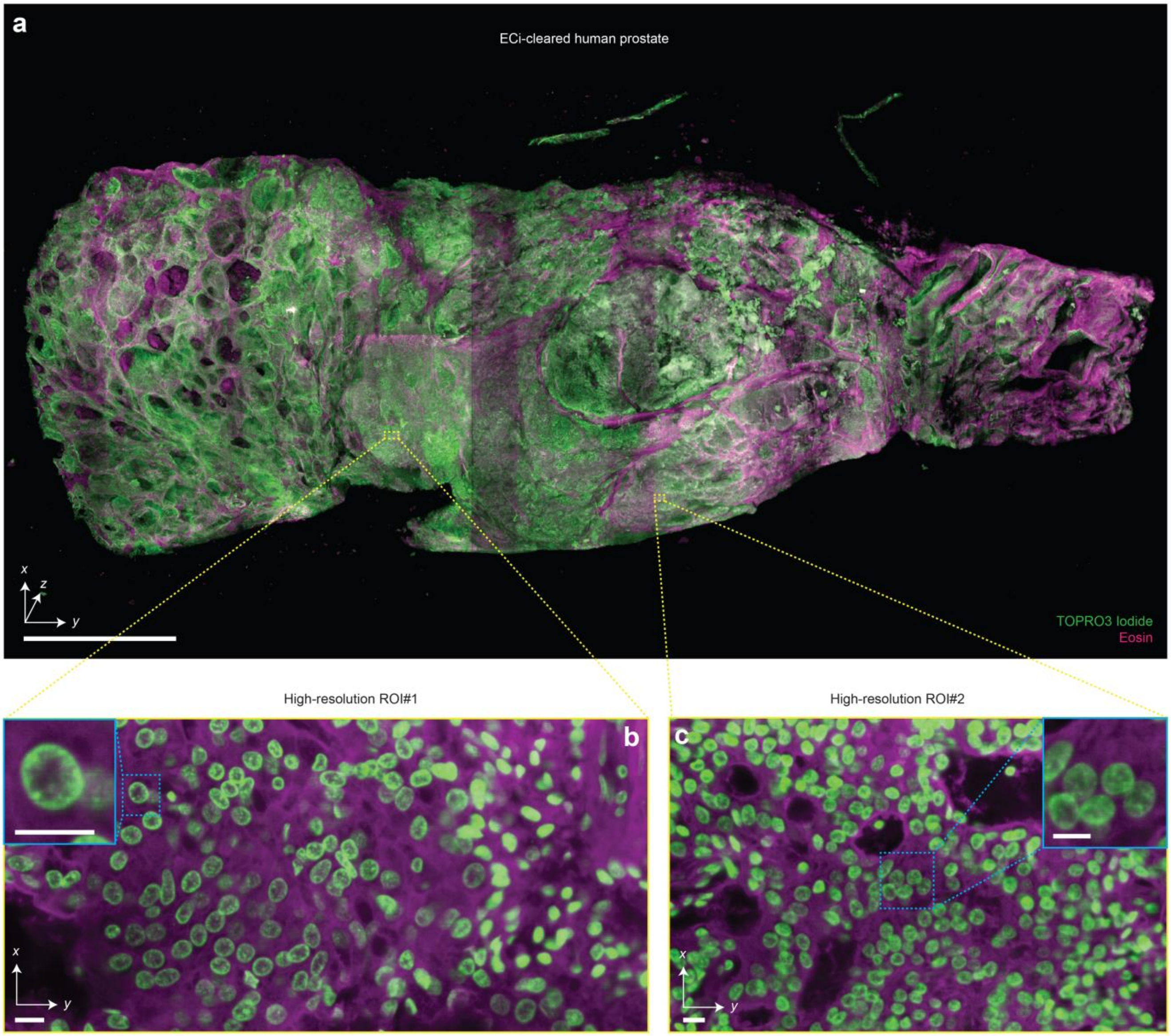


**Supplementary Figure 18 | Multi-scale 3D pathology of human prostate tissue. (a)** Fast meso-scale screening results for a multi-cm-sized piece of human prostate tissue stained with TO-PRO-3 Iodide (nuclear) and Eosin. Representative high-resolution regions of interests for two different foci of cancer are shown in **(b)** and **(c)**. The insets demonstrate the ability to clearly resolve sub-nuclear features in cancer nuclei. Scale bar lengths are as follows: **(a)** 1 cm, **(b)** and **(c)** 10 μm.

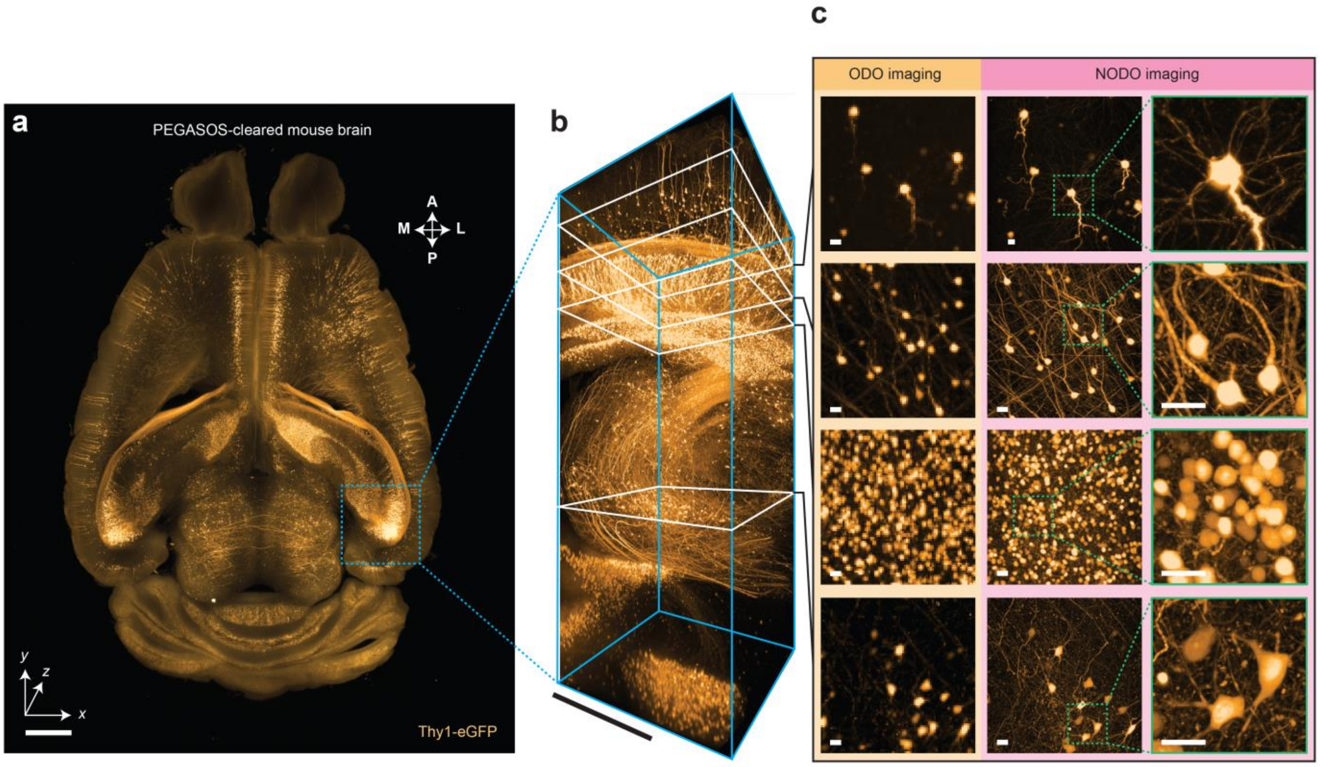


**Supplementary Figure 19 | Multi-scale interrogation of PEGASOS-cleared mouse brain. (a)** Whole brain ODO imaging of a Thy1-eGFP mouse brain cleared using the PEGASOS protocol. **(b)** Region of interest z-stack imaged using NODO imaging. Individual z-planes are shown in **(c).** Matched z-planes from the ODO imaging dataset are also shown for comparative purposes. Scale bar lengths are as follows: **(a)** 1 mm, **(b)** 0.5 mm, and **(c)** 10 μm.

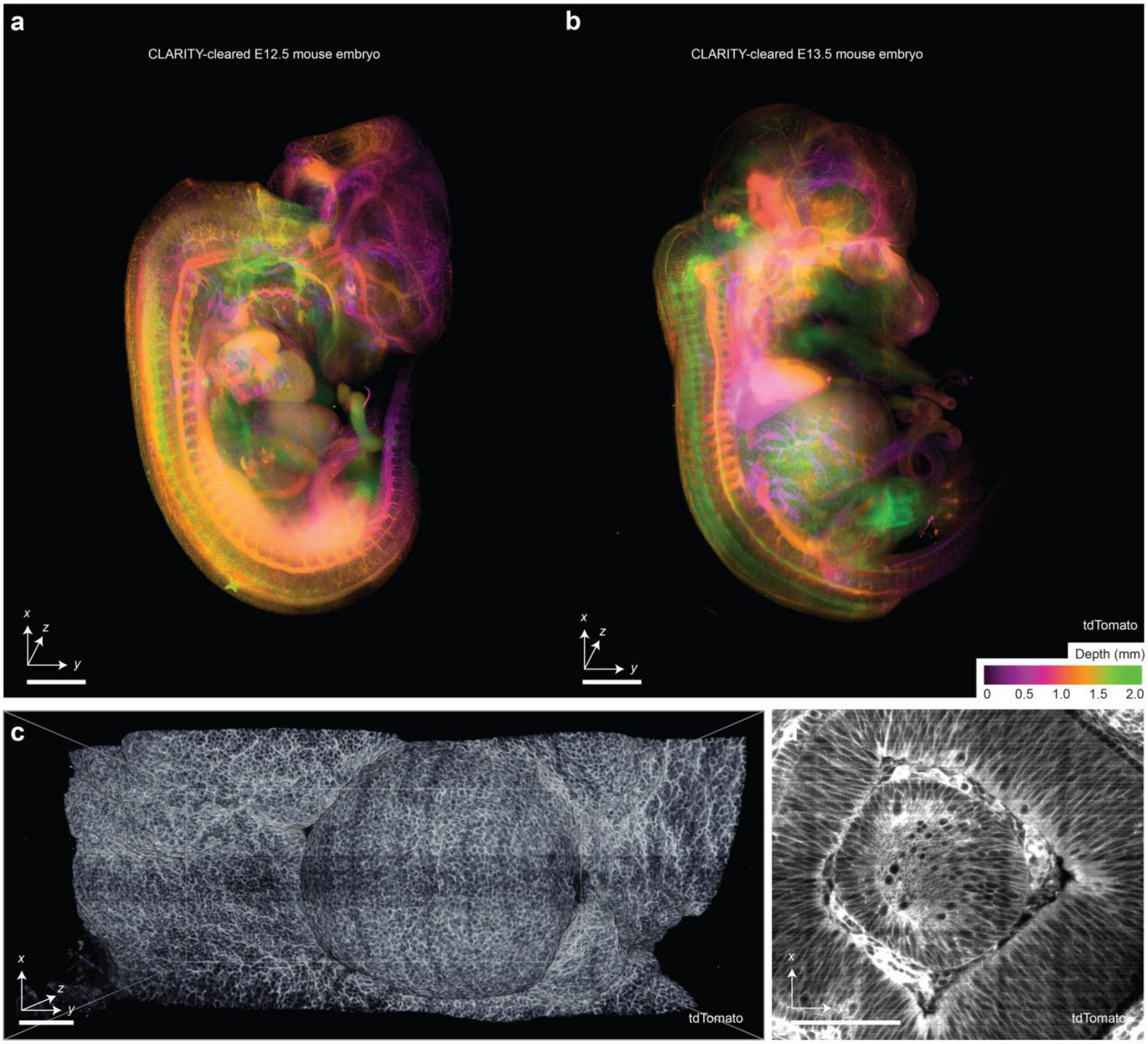


**Supplementary Figure 20 | Mid-gestation mouse embryos reporting cell boundaries using SHIELD**. **(a)** Intact embryonic day (E)12.5 and **(b)** E13.5 mouse embryo with endogenous membrane-tagged tdTomato fluorescence signal indicating cell borders. Embryos are depth-coded in *z*. A higher-resolution region of interest focused on the eye of **(a)** is shown in **(c)** and **(d)**. An additional zoom-in highlights the ability to resolve diversity of cell boundary shapes **(d)**. Scale bars lengths are as follows: **(a)** 1 mm, **(b)** and **(c)** 100 μm.

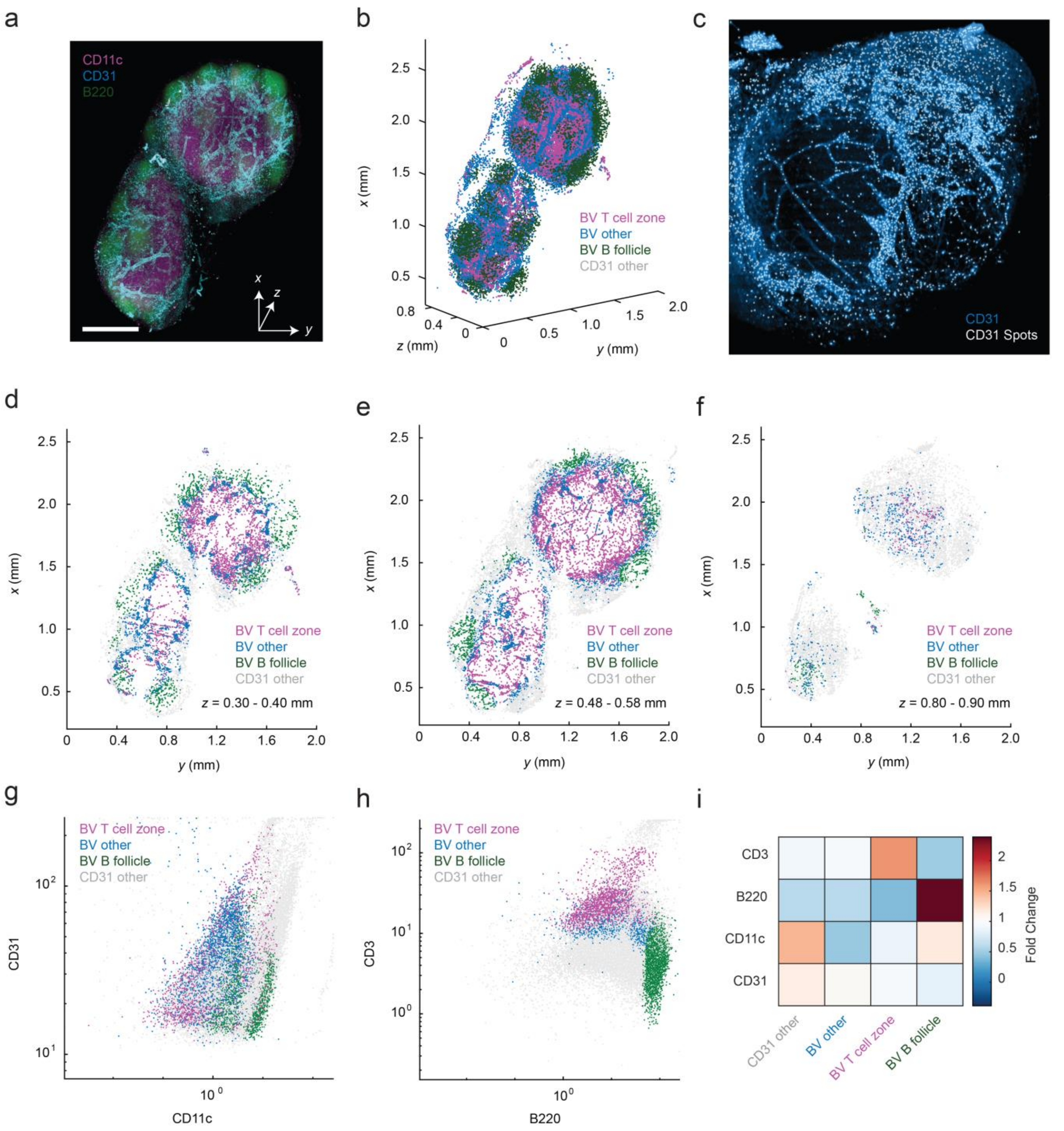


**Supplementary Figure 21 | CytoMAP analysis of CD31 signal in Ce3D-cleared mouse lymph node. (a)** ODO imaging of a whole Ly5.1 mouse lymph node cleared using the Ce3D protocol. **(b)** 3D plot of CD31-positive spots (i.e. central coordinates of an object) color-coded by cluster type as defined by CytoMAP. **(c)** Cross section of the cleared mouse lymph node with CD31 spots overlaid on the CD31 fluorescence channel showing sampling of the positions of vasculature and other CD31-positive objects. **(d-f)** 3D point plots of the CD31 spots color coded by cluster type from 0.1-mm thick cross sections from different *z* positions of the lymph node showing the distinct spatial patterns of the different CD31 clusters. **(g-h)** Plots of the mean fluorescent intensity of the CD31 spots color coded by cluster type. **(i)** Heatmap of the fold change of the mean fluorescent intensity of the CD31 spots showing the expression profile of the spots from individual clusters. **(g-i)** Expression profiles of CD31 spots, including signals from CD31-expressing cells and most proximal neighbors.

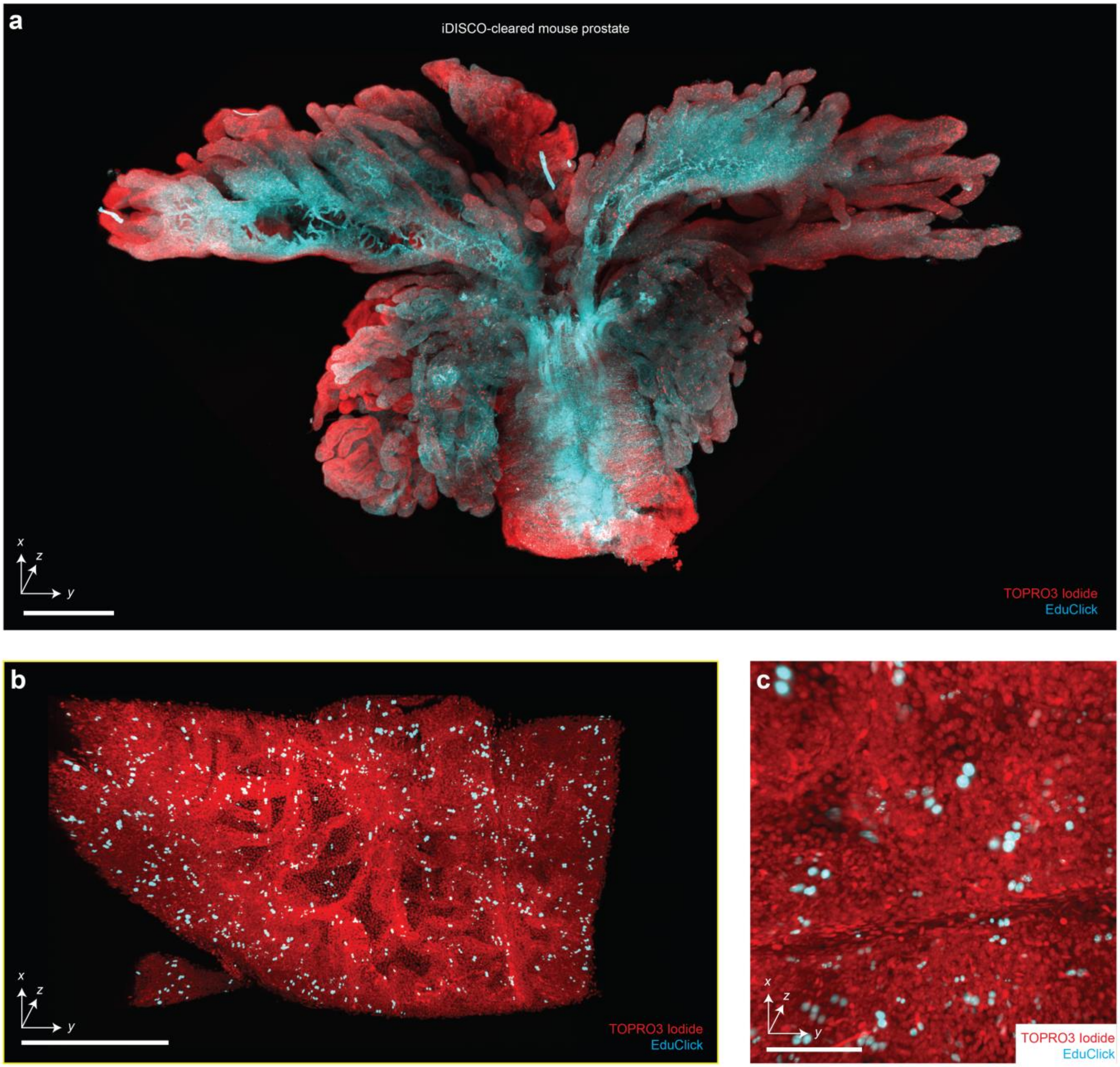


**Supplementary Figure 22 | Assessment of 3D cell proliferation assays with iDISCO. (a)** Screening of an intact mouse prostate cleared using iDISCO, labeled with TOPRO3 Iodide (nuclear) and an EduClick cell-proliferation marker. A higher-resolution region of interest focused on a prostate gland is shown in **(b)**. An additional zoomed-in view showing the ability to resolve individual proliferating nuclei **(c)**. Scale bars lengths are as follows: **(a)** 1 mm, **(b)** 200 μm, and **(c)** 100 μm.

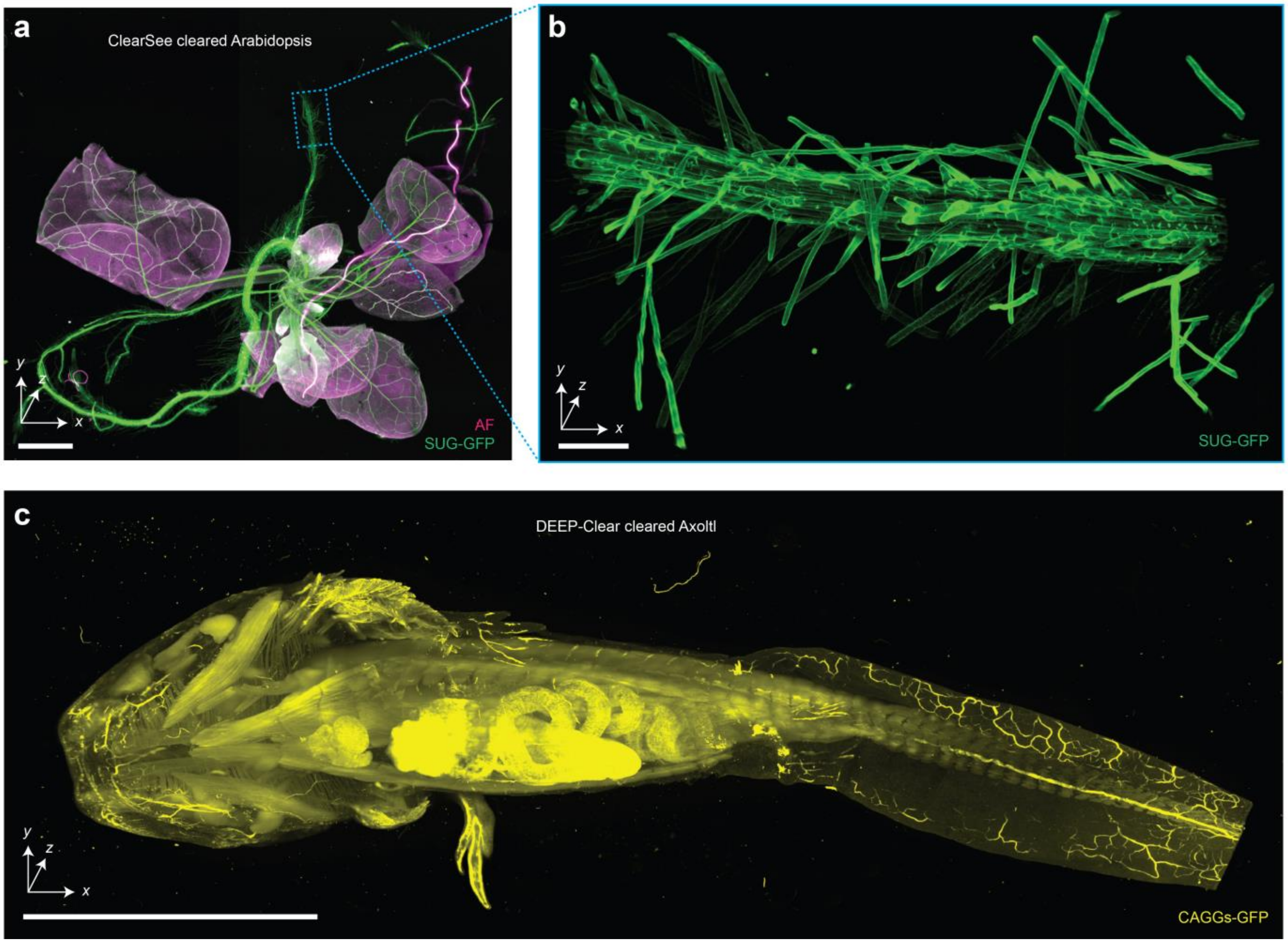


**Supplementary Figure 23 | Imaging of non-rodent and non-human tissues. (a)** Hybrid open-top light-sheet imaging of a ClearSee-cleared Arabidopsis specimen. **(b)** Higher-resolution imaging of the Arabidopsis root. **(c)** Meso-scale imaging of a large multi-cm Axolotl cleared with DEEP-Clear. Scale bars lengths are as follows: **(a)** 1 mm, **(b)** 100 µm, and **(c)** 1 cm.

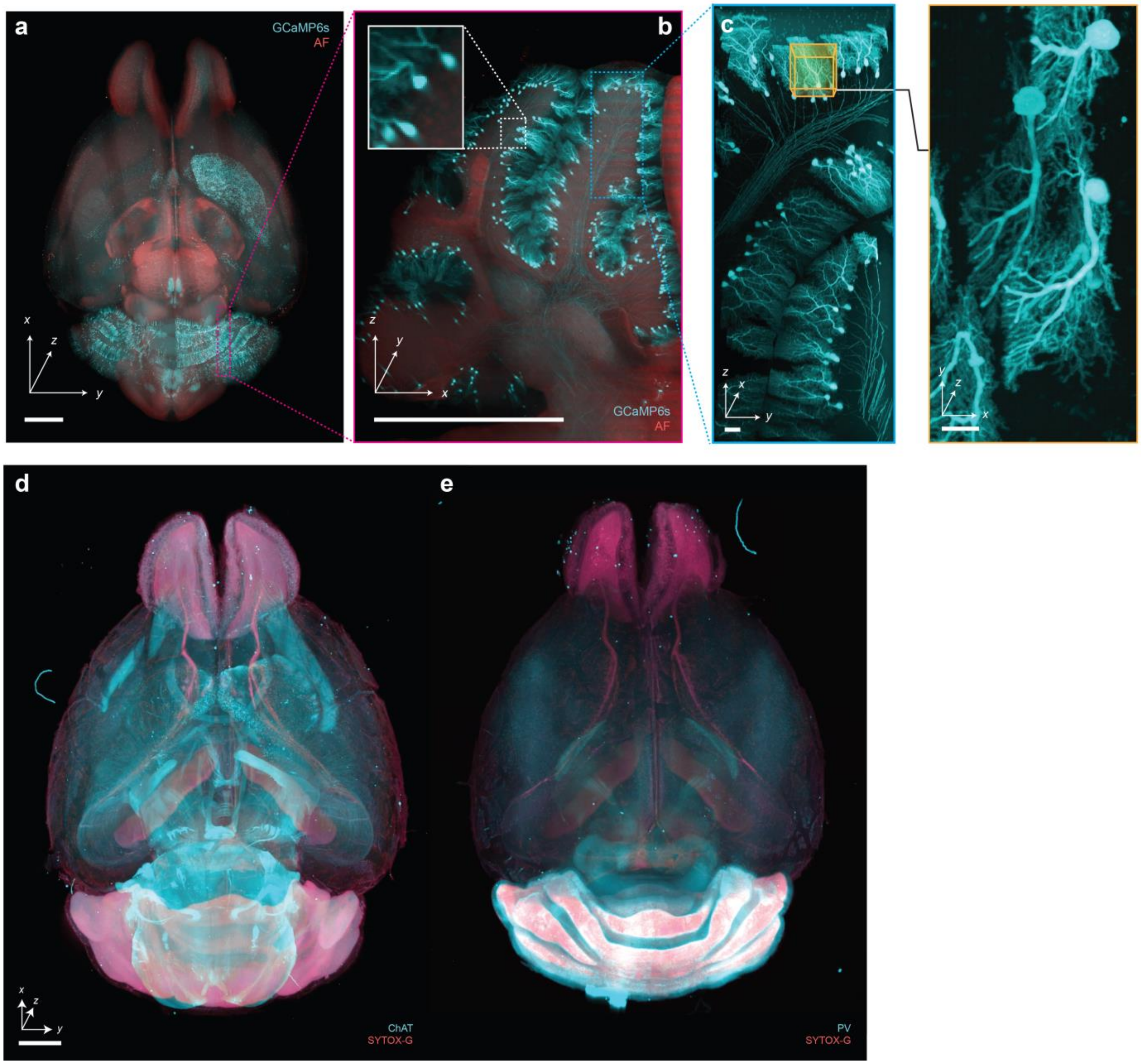


**Supplementary Figure 24 | Imaging of immunostained and endogenously fluorescent CUBIC-cleared mouse brains.** Hybrid OTLS imaging of whole mouse brains. **(a-c)** with endogenously preserved GCaMP6s fluorescence, and immunostained with **(d)** anti-ChAT antibody + SYTOX-G or **(e)** anti-Parvalbumin (PV) antibody + SYTOX-G by CUBIC-HistoVision. Scale bar lengths are as follows: **(a-b)** 1 mm, **(c)** 10 μm, **(d-e)** 2 mm.

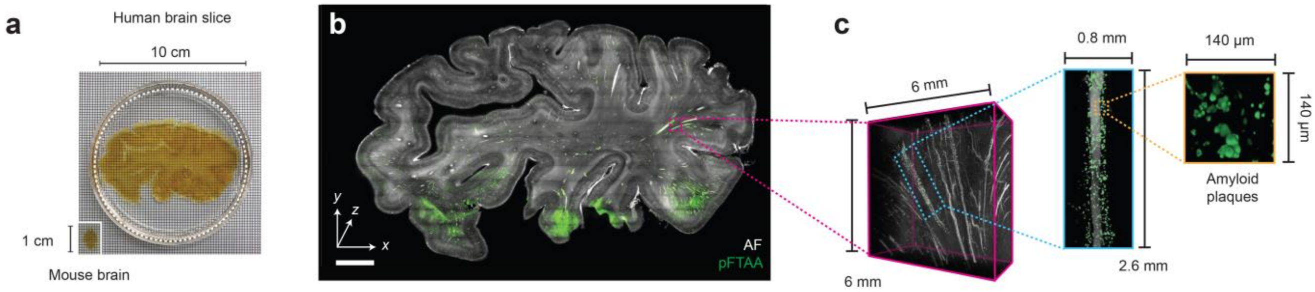


**Supplementary Figure 25 | Large-scale imaging of thick human tissues. (a)** Slab of human brain tissue after CUBIC clearing. A mouse brain is shown for size comparison. **(b)** ODO imaging results of the entire 3-mm thick brain slab. Autofluorescence is shown in black and white, and the amyloid small molecule stain (pFTAA) is shown in green. **(c)** Zoom-in views of an amyloid-rich region reveal perivascular accumulation. Scale bar lengths are as follows: **(b)** 1 cm.

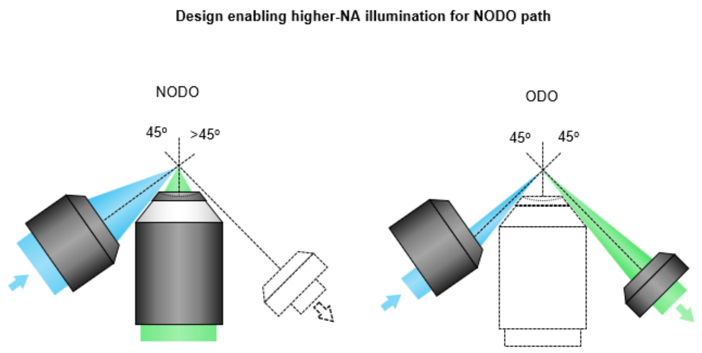


**Supplementary Figure 26 | Higher-NA illumination for NODO path.** To improve the axial resolution of the NODO path, a higher illumination NA could be used where the central axis of the objective is >45 deg with respect to the NODO collection objective. ODO imaging is still possible by only illuminating the edge of the illumination objective's pupil.

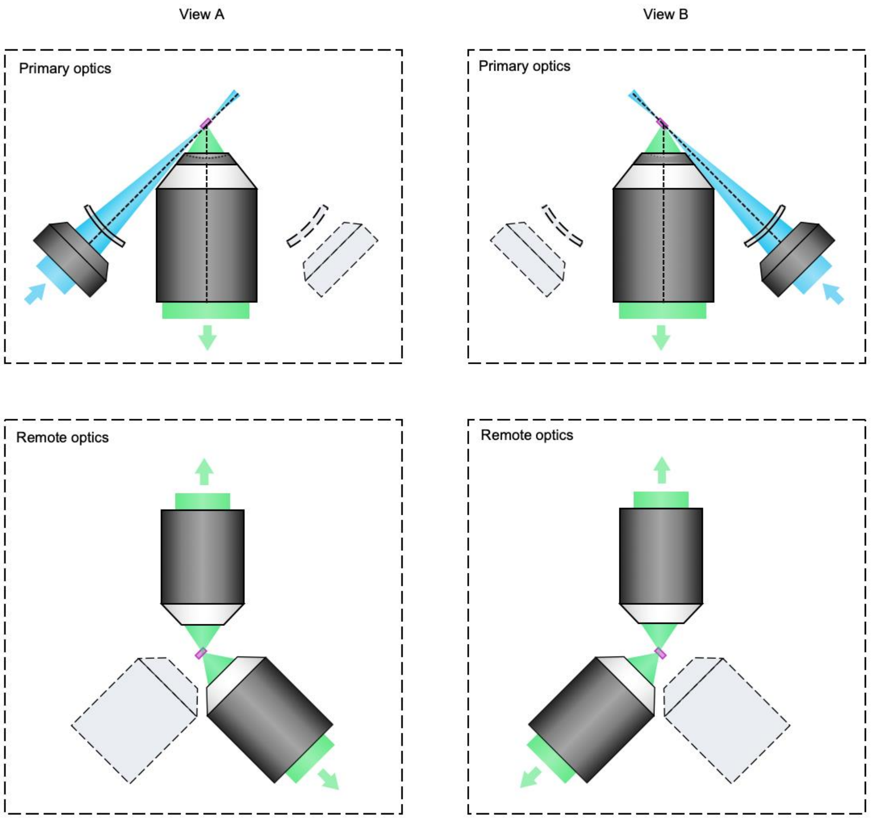


**Supplementary Figure 27 | Dual-view hybrid OTLS system.** Although in the current hybrid system, only one of the angled optical paths is used for illumination, and the other for ODO imaging, in the future both paths could be used for illumination and collection. As shown, this would yield two views (A and B) of the specimen, which could be input into a fusion deconvolution algorithm to achieve more isotropic resolution for both NODO imaging paths. It is important to note that it would be challenging to accommodate two remote objectives for single-objective light-sheet microscope designs due to the increased tilt angle at the remote focus, which often necessitates bespoke objectives with unique geometric constraints. However, our NODO design reduces the required tilt angle, and therefore may permit the use of two angled air objectives (off-the-shelf) at the remote focus. Note dual-view imaging could also be achieved by using two illumination paths but a single remote objective, as detailed in [18, 37].

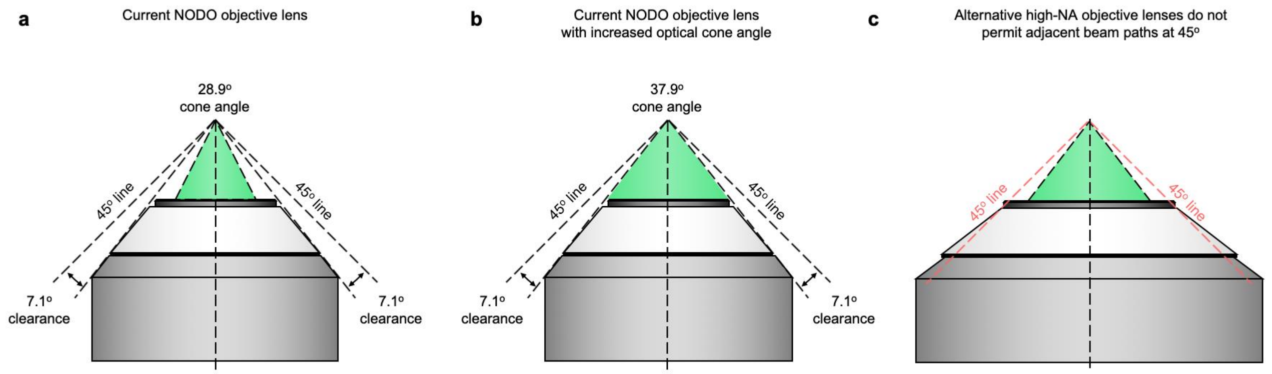


**Supplementary Figure 28 | Considerations for NODO objective. (a)** Schematic of the current NODO objective. The mechanical housing of the objective permits optical paths at 45 deg on both sides, with a maximum cone angle of 7.1 deg (NA ~ 0.12 in air). **(b)** The current NODO objective leaves ~ 9 deg between the optical and mechanical cone angles. Theoretically, the optical cone angle could be expanded from 28.9 to 37.9 deg in a future customized multi-immersion objective. In practice, this angle would be slightly less when accounting for the objective's finite FOV. **(c)** The majority of alternative high-NA immersion objectives (e.g., NA ~ 1.0) have mechanical housings that do not permit optical paths at 45 deg for the addition of ODO imaging.

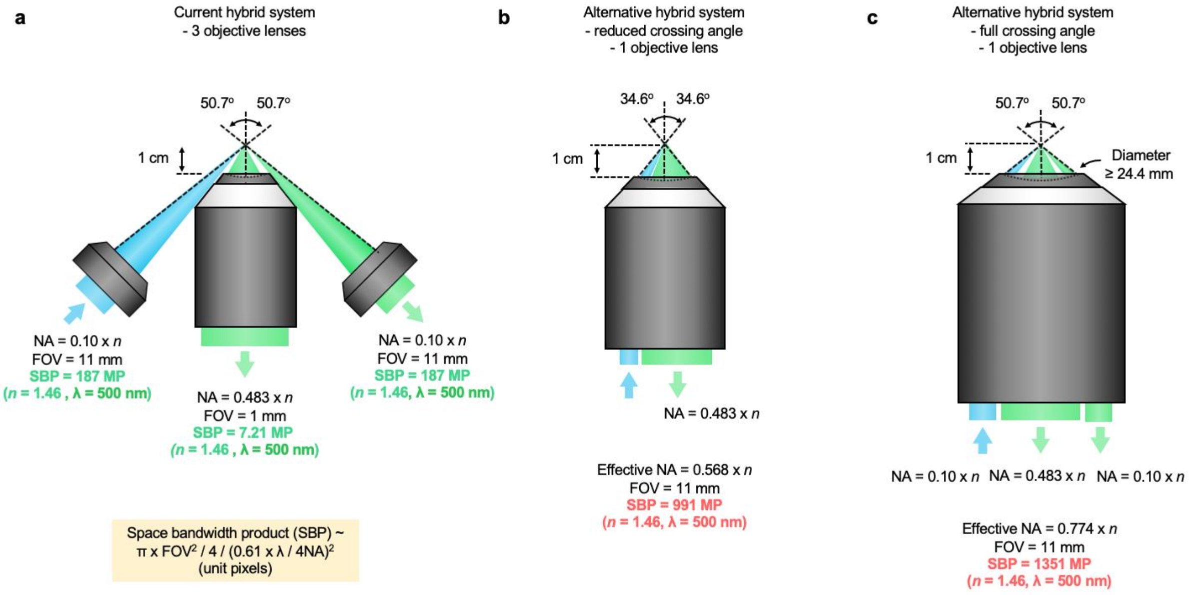


**Supplementary Figure 29 | Alternative designs based on customized objectives. (a)** The current hybrid OTLS system uses three objectives for illumination, high-resolution NODO imaging, and low-resolution ODO imaging. These objectives are all commercially available. **(b)** In theory, all three optical paths could be incorporated into a single objective. This theoretical objective would require an effective NA of 0.568 (in air) and a large 11 mm field of view (FOV). This NA would reduce the crossing angle of the illumination and collection beam paths but provide identical NAs to the current system. However, these specifications are extremely demanding, and would require a space bandwidth product (SBP) of ~ 991 megapixels (MP), which is far beyond any currently produced objectives. **(c)** Another alternative design would maintain the same crossing angles as the current system, instead delivering and collecting all beam paths through a single massive objective. The specifications for this objective would be even more demanding, with SBP ~ 1351 MP. These alternative designs are highly speculative, and it is not likely that these objectives would be technically or financially feasible to design and/or manufacture.

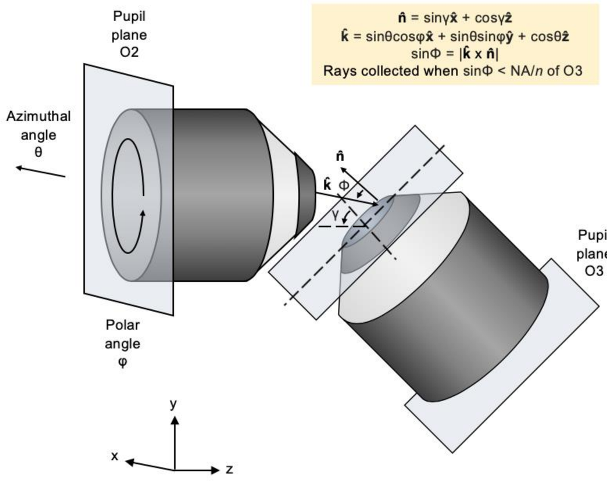


**Supplementary Figure 30 | Numerical simulation geometry.** Based on a spherical coordinate system, each ray exiting objective O2 can be described by an azimuthal angle, $\theta$, and polar angle, $\varphi$. The direction of that light ray is then described as $\hat{k} = \sin\theta\cos\varphi\hat{x} + \sin\theta\sin\varphi\hat{y} + \cos\theta\hat{z}$. The surface normal from objective O3 is described by $\hat{n} = \sin\gamma\hat{x} + \cos\gamma\hat{z}$. Therefore, the angle of incidence of the ray on O3 is given by $\sin\Phi = |\hat{k} \times \hat{n}|$. The incident pupil plane from O2 can then be clipped by the NA of O3.

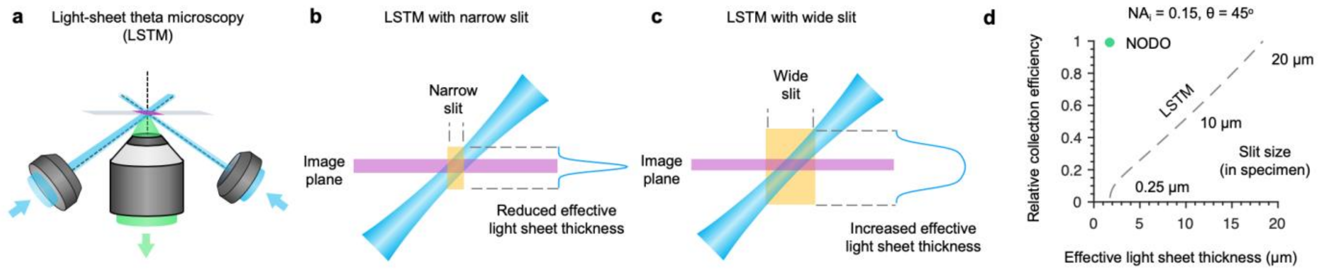


**Supplementary Figure 31 | Comparison of light-collection efficiency and effective light-sheet thickness for LSTM and NODO architectures. (a)** Rather than using a remote focus to re-image a tilted light sheet, LSTM laterally scans two illuminations beams to generate a virtual light sheet within the specimen that is orthogonal to the collection objective. **(b-c)** To reject out-of-focus light, the scanning beams are synced to a confocal slit. A narrow slit maintains a thin effective light-sheet thickness, at the cost of reduced light-collection efficiency. While using a wider slit would increase light efficiency, it would also increase the effective light-sheet thickness due to imaging regions of the light sheet that are outside the image plane (i.e., the focal plane of the collection objective). For illustrative purposes, only one of the two illumination beams is shown. **(d)** Simulations were performed comparing NODO and LSTM architectures using the specifications of the current NODO system ($NA_i$ = 0.15 and θ = 45 deg). For the same effective light-sheet thickness, the light-collection efficiency for LSTM is <10% compared to NODO. Likewise, to achieve the same light-collection efficiency as NODO, the effective light-sheet thickness for LSTM would be increased from 1.9 µm to 18.4 µm. Therefore, light-collection efficiency and effective light-sheet thickness are inherently coupled for LSTM, whereas they are decoupled for NODO. Note, for NODO imaging, the sCMOS camera is adjusted to match the confocal parameter ($2z_R$) of the Gaussian light sheet, which is ~20 µm.
